# Moran Process Version of the Tug-of-War Model: Complex Behavior Revealed by Mathematical Analysis and Simulation Studies

**DOI:** 10.1101/2021.10.20.465201

**Authors:** Adam Bobrowski, Marek Kimmel, Monika K. Kurpas, Elżbieta Ratajczyk

## Abstract

In a series of publications McFarland and co-authors introduced the tug-of-war model of evolution of cancer cell populations. The model is explaining the joint effect of rare advantageous and frequent slightly deleterious mutations, which may be identifiable with driver and passenger mutations in cancer. In this paper, we put the Tug-of-War model in the framework of a denumerable-type Moran process and use mathematics and simulations to understand its behavior. The model is associated with a time-continuous Markov Chain (MC), with a generator that can be split into a sum of the drift and selection process part and of the mutation process part. Operator semigroup theory is then employed to prove that the MC does not explode, as well as to characterize a strong-drift limit version of the MC which displays “instant fixation” effect, which was an assumption in the original McFarland’s model. Mathematical results are fully confirmed by simulations of the complete and limit versions. They also visualize complex stochastic transients and genealogies of clones arising in the model.

## 1. Introduction

The Tug-of-War model was developed in a series of papers of McFarland and co-authors [21, 23, 24] to account for existence of mutually counteracting rare advantageous driver mutations and more frequent slightly deleterious passenger mutations in cancer. In its original version it is a state-dependent branching process, analyzed by a range of simulation methods and analytical approximations.

We adopt a different, simpler, approach, in which we reformulate McFarland’s original definition to put it into the framework of a Moran model, which we investigate by complementary methods of mathematical analysis and simulation.

In the current study we are not primarily concerned with understanding the genealogies of the individuals such as cancer cells present in the populations. We identify individuals with the same counts of passenger and driver mutations and follow trajectories of the so-defined types. As it will become clear in the sequel, process behavior is quite complicated. Nevertheless, we demonstrate absorption properties of the process with no mutations (Section 4) and use operator semigroup theory to prove two limit cases (Section 7).

## 2. The model: a population under selection, drift and mutation

We consider a population of a fixed number *N* of individuals, each of them characterized by a pair of integers (*α, β*), corresponding to the numbers of drivers and passengers in its genotype, respectively. This pair determines the fitness *f* of the individual by the formula

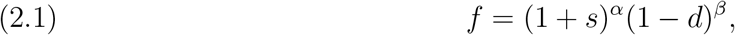

where *s* > 0 and *d* ∈ (0, 1) are certain parameters describing selective advantage of driver mutations over passenger mutations. Thus, the entire population may be identified with the vector

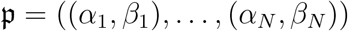

of *N* pairs of integers, with the accompanying vector

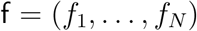

of fitnesses.

The population is under drift and selection pressure: the individual of type (*α_i_, β_i_*) lives for an exponential time with parameter

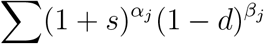

where the sum is over all *j* = 1,…, *N* such that (*α_j_, β_j_*) ≠ (*α_i_, β_i_*), and then is replaced by an individual of different type. More specifically, let *n_α,β_* be the number of individuals of type (*α, β*) and *n* be the number of different types of individuals in the population, then the time to the death of each individual of type (*α_i_, β_i_*) is the minium of *n* – 1 exponential random variables *T_α_j_, β_j__* where (*α_j_, β_j_*) ≠ (*α_i_, β_i_*) and *T_α_j_, β_j__* has parameter *n_α_j_, β_j__*(1 + *s*)^*α_j_*^(1 – *d*)^*β_j_*^. Upon this individual’s death, conditional on the minimal time being equal to *T_α_k_,α_k__*, this individual is replaced by one of the individuals of type (*α_k_, β_k_*), each if these individuals being equally likely. This process then continues with 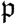 modified by replacing its *i*th coordinate (*α_i_, β_i_*) by its *k*th coordinate (*α_k_, β_k_*).

In particular, if all individuals in 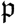 are pairwise different, the time to the first drift and selection event for the entire population is exponential with parameter (*N* – 1)Σf where

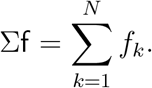

After this time is over, one individual dies and is replaced by an exact copy of one of the remaining individuals, the probability that the *i*th individual dies and is replaced by the jth (*j* ≠ *i*) being 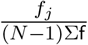. If, on the other hand, all individuals are the same, nothing happens: there are no drift and selection events.

Moreover, each individual may, after an independent exponential time with parameter, say λ, and independently of other individuals, undergo a mutation event, changing its state to either (*α* + 1, *β*) or (*α, β* + 1) with (conditional) probabilities *p* ∈ (0,1) and *q*, respectively. In other words, all mutations occur at the epochs of a Poisson process with intensity λ, occurrences of driver mutations on each individual form a colored Poisson process, with probability of coloring equal p, and the occurrences of passenger mutations form a colored Poisson process with probability of coloring equal *q*. It follows (see the Colouring Theorem on p. 53 in [17]) that driver and passenger mutations form Poisson processes with parameters

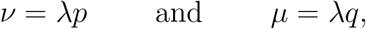

respectively, and these processes are independent of each other, and independent of mutation processes on other individuals. (Although technically we never use this assumption, what we have in mind is the case where *p* is significantly smaller than 1 – *p*, so that long strings of passenger mutations are interrupted by rare driver mutations.) In particular, given that initially an individual’s fitness is *f*, after time *t* its expected fitness is the product of *f*, e^−λ*pt*^e^λ*p*(1+*s*)*t*^ (the contribution of driver mutations) and e^−λ*qt*^e^λ*q*(1−*d*)*t*^ (the contribution of passenger mutations), and thus equals

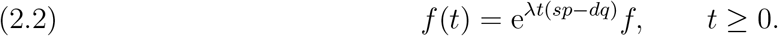

In a number of cells this expected fitness does not grow to infinity or decay to zero; such cells are thus characterized by the following balance condition for the introduced parameters:

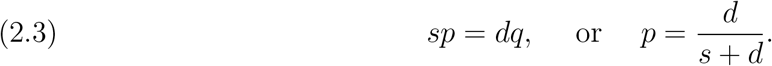

In other words, the advantage gained by a driver mutation is balanced by the small probability of such event.

In other cells, however, driver mutations, though rare may have a slight edge over the passenger mutations caused by large *s*. Such cell populations are characterized by

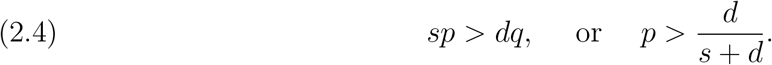

In yet different populations, driver mutations will be so rare that the expected total fitness diminishes in time. To characterize such populations, we reverse the inequalities in (2.4).

## 3. A Markov chain and the related intensity matrix

The population described in Section 2 is modeled by a stochastic process

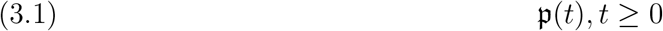

with values in the state-space 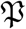 of *N* ordered copies of the Cartesian product 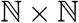, where 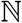 is the set of natural numbers:

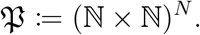

This is just to say that at each time *t*, the population is an *N*-tuple of pairs (*α_i_*(*t*), *β_i_*(*t*)), 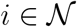 of positive integers, where

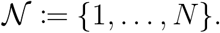

Since 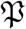 is a countable set, the process 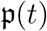, *t* ≥ 0 may be thought of as a time-continuous Markov chain.

Such Markov chains are conveniently described by means of intensity (Kolmogorov) matrices that gather information on rates (intensities) with which these processes leave a given state and jump to other states (see e.g. [4, 25]; see also our Section 10). We will write the intensity matrix for the process (3.1) as the sum of two intensity matrices representing mutations and drift and selection events, respectively.

To describe the first of these, call it *Q_M_*, (‘*M*’ for ‘mutations’) let *D* and *P* (‘*D*’ for driver and ‘*P*’ for passenger) be the following maps of 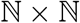 into itself:

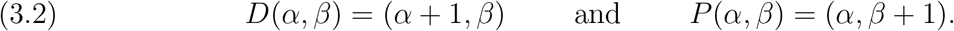

Moreover, for each *i* = 1,…, *N*, let 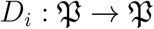 be the map in which the *i*th coordinate (*α_i_, β_i_*) of a 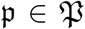 is replaced by *D*(*α_i_, β_i_*). Similarly, let 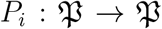 be the map in which the *i*th coordinate (*α_i_, β_i_*) of a 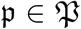 is replaced by *P*(*α_i_, β_i_*). In these notations, the intensity 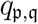 of going from a state 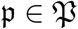 to a state 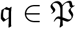 in the mutation process is

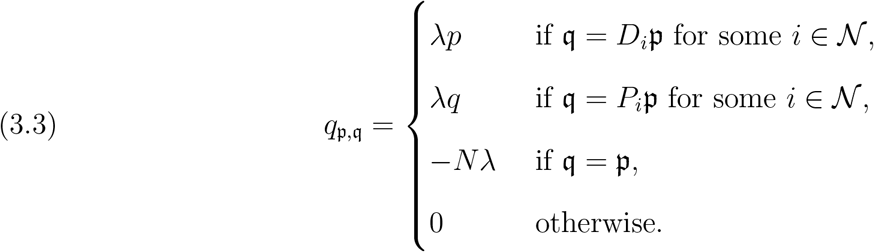

Similarly, for *i, j* = 1,…, *N* let 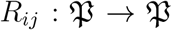 be the map that replaces the *i*th coordinate (*α_i_, β_i_*) of a 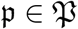 by its jth coordinate (*α_j_, β_j_*), leaving the remaining coordinates intact. For example, if *N* = 3, *R*_1,3_ maps ((*α*_1_, *β*_1_), (*α*_2_, *β*_2_), (*α*_3_, *β*_3_)) to ((*α*_3_, *β*_3_), (*α*_2_, *β*_2_), (*α*_3_, *β*_3_)). The intensity matrix describing drift and selection events, say *Q_S_*, has then the following entries:

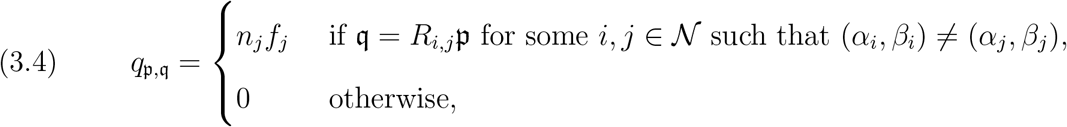

where *n_j_* is the number of individuals in 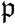 that are identical to the individual number *j* and 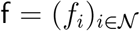 is the vector of fitnesses of individuals in 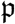. More specifically,

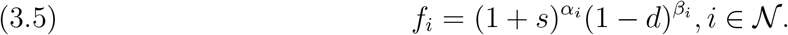

This formula does not cover the case where 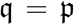 because this case requires a bit of preparation. Namely, let *n_α,β_* denote the number of individuals of type (*α, β*) so that in particular 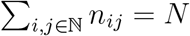. Then,

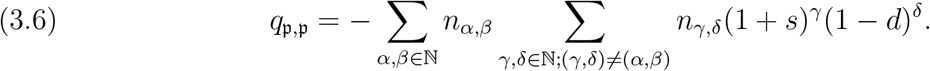

We note that in the case where all individuals in a population 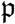 are different, the formula for 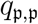 simplifies to:

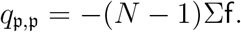

On the other hand, if all individuals in this population are of the same type, 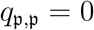.

Finally, the entries in the intensity matrix *Q* for the entire chain (3.1) are sums of the entries of matrices *Q_M_* and *Q_S_*:

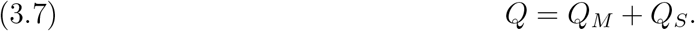

## 4. Properties of the drift and selection chain

Consider the evolution of a population when mutations are absent, and only drift and selection events, as described above, are possible. This evolution is governed the intensity matrix *Q_S_* with entries given in (3.4) and (3.6).

For definiteness, let 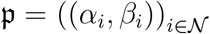 be the initial state of this population and assume that all its individuals have different characteristics, i.e. (*α_i_, β_i_*) ≠ (*α_j_, β_j_*) for *i* ≠ *j*. It is rather easy to see first of all there is only a finite number of states that can be reached from the state 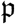: more precisely, there are at most *N_N_* such states (including 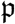 itself). For, since the chain is that of replacing coordinates of 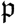 by copies of other coordinates, there are only *N* possibilities for the first coordinate of future states, *N* possibilities for the second coordinate, and so on.

Second, all these states, except for those with all coordinates equal, i.e. except for

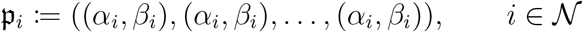

are transient for this chain. Indeed, for any other state there is a non-zero probability that the number of different individuals in the population will decrease in the next drift and selection event. Since the rules of the chain do not allow jumps from the states with smaller number of different individuals to the states with larger number of different individuals, the process will never come back to the state under consideration. This shows that this state cannot be recurrent, and thus, by the well-known dichotomy (see e.g. [25], Section 3.4) it must be transient. On the other hand, all the states 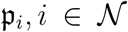 are absorbing. Hence, the process starting at a 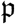 must eventually end up at one of 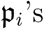. Certainly, in the case where not all individuals in the original population are different, the fate of the population is similar: it is only the number of possibilities for the end population that is smaller. We summarize our discussion in the following theorem.

### Theorem 4.1.

*Let* 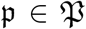 *be a population and let M be the number of different variants in* 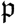. *Then, there is a set* 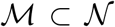 *such that (a)* 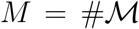, *and (b)* 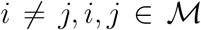 *implies* (*α_i_, β_i_*) = (*α_j_, β_j_*). *For* 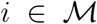 *let* 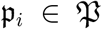 *be the population in which all individuals are identical to each other and to the *i*th individual in the original population* 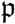. *Then, the drift and selection chain starting at* 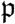 *will eventually end up at* 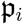 *with certain probability* 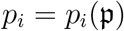 *where* 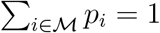.

We note in passing that whereas there could be many choices of 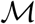, the thesis of our theorem remains the same for all of them.

This theorem is a reflection of the fact that drift and selection chain strives to reduce the number of variants in the population by removing randomly selected variants and replacing them by other variants; in the absence of other forces, and mutation in particular, the chain’s operation in the long run leads to fixation of one the variants. What this theorem does not express openly is that drift and selection chain favors variants with larger fitness. The latter information, besides being visible in formula (3.4), is hidden in the probabilities p_i_ featuring in Theorem 4.1; roughly speaking, the larger the fitness of an individual, the larger is the probability of fixation of its variant. Notably, even though selection favors variants with larger fitness, it acts together with genetic drift which may ‘blindly’, by chance, remove better fit variants from the population. Hence, the fact that a variant with larger fitness is favored by selection results in a higher probability of its fixation, and not in the inevitability of its fixation.

In what follows we will see this principle expressed in explicit formulae for *p_i_*’s in the cases *N* = 2 and *N* = 3, considered here for the sake of illustration. In our calculations it will be convenient, for the sake of shortening our equations and making figures readable, to identify a population 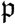, which is an *N*-dimensional vector of pairs of integers, with the *N*-dimensional vector of corresponding fitnesses calculated by formula (3.5), the latter vector being half as long as the former. Although it is possible, by an appropriate choice of parameters, to have two different individuals with the same fitnesses, i.e. to have (*α_i_, β_i_*) ≠ (*α_j_, β_j_*) and at the same time (1 + *s*)^*α_i_*^ (1 – *d*)^*β_i_*^ = (1 + *s*)^*α_j_*^(1 – *d*)^*β_j_*^, such an identification should not lead to misunderstandings.

For *N* = 2 the probabilities *p*_1_ and *p*_2_ are easily calculated explicitly: unless it is already uniform, a population 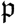 with fitness (*f*_1_, *f*_2_), after an exponential time with parameter *f*_1_ + *f*_2_, becomes 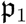 with probability 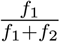 or 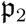 with probability 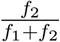. This shows that 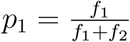 and 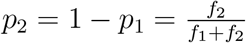.

Somewhat surprisingly, already for *N* = 3 the formulae for *p_i_*’s are more complicated, and do not follow the perhaps expected pattern 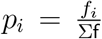. Before we see that, however, we note the following important property of the chain under consideration: Let us call two states f = (*f*_1_,…, *f_N_*) and 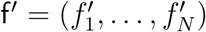 *associated* if there is a permutation Π of the set 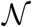 such that 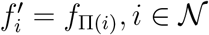. The property we want to note is as follows.

### Theorem 4.2.

*The drift and selection processes starting at two associated states are analogous*.

What we mean by that proposition is that (a) the times to the first drift and selection events for either of two associated states f and f′ have the same distribution, (b) the probability that in such an event the *i*th coordinate of f is replaced by the *j*th, is the same as the probability that the Π(i)th coordinate of ***f*** is replaced by the Π(*j*)th, and (c) if in these drift and selection events the *i*th coordinate of f is replaced by the *j*th, and the Π(*i*)th coordinate of f′ is replaced by the Π(*j*)th then the states after these events are again associated. These statements are clear from the description of the drift and selection chain, and combined together prove Theorem 4.2.

We are now ready to find *p_i_*s for *N* = 3. We think of a process that starts at an f = (*f*_1_, *f*_2_, *f*_3_). Figure 1 illustrates the fact that in order to reach the state f_1_ = (*f*_1_, *f*_1_, *f*_1_) this process must go through (*f*_1_, *f*_1_, *f*_3_), (*f*_1_, *f*_2_, *f*_1_) or one of their associates. The first of these states is reached directly with probability 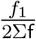. This state or one of its associates may also be reached indirectly, via (*f*_1_, *f*_3_, *f*_3_), with probability 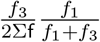. Thus, the probability of reaching (*f*_1_, *f*_1_, *f*_3_) or one of its associates is

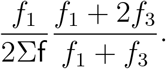

**Figure 1.**
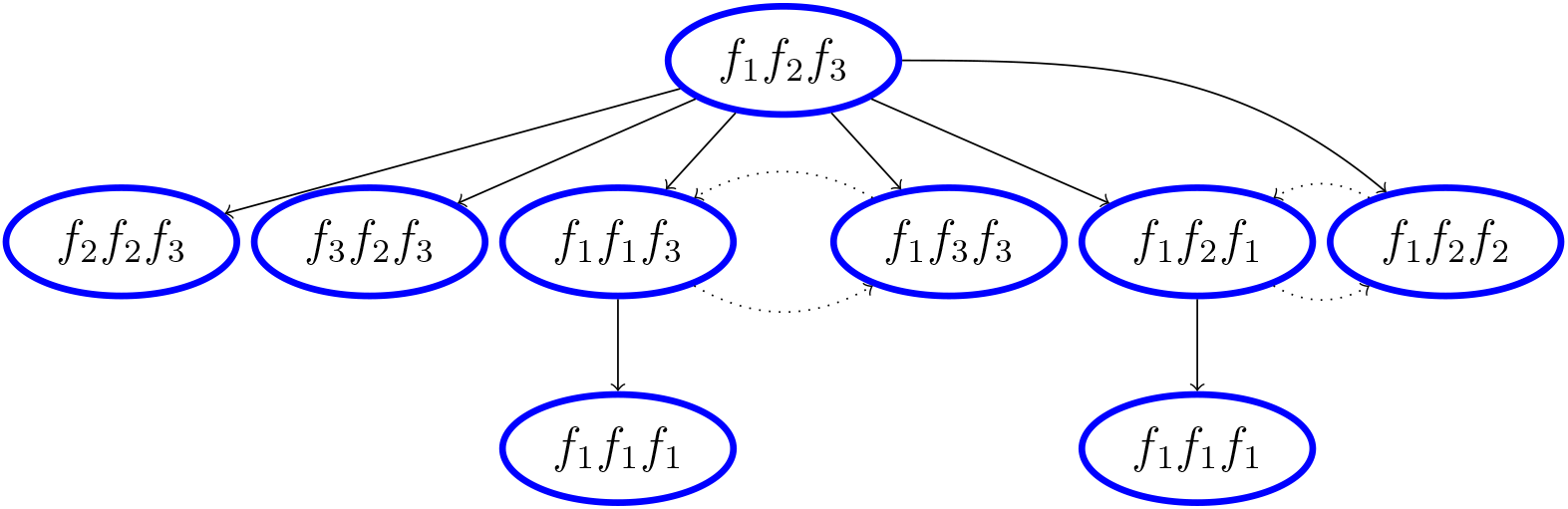
Calculating probability *p*_1_ in the case *N* = 3. Dotted lines denote communication between events associated with (*f*_1_, *f*_1_, *f*_3_) and (*f*_1_, *f*_3_, *f*_3_), and (*f*_1_, *f*_2_, *f*_1_) and (*f*_1_, *f*_2_, *f*_2_).

Then, before reaching f_1_ from one of these associated states, the process may visit an associate of (*f*_1_, *f*_3_, *f*_3_), and this may happen *k* ≥ 0 times. Since the properties of the processes starting from associated states are analogous, the probability of reaching f_1_ from one of associates of (*f*_1_, *f*_1_, *f*_3_) is

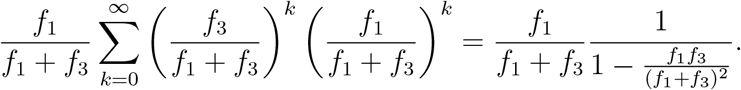

Therefore, the probability that f_1_ will be reached through (*f*_1_, *f*_1_, *f*_3_) or its associate is 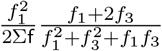 and so

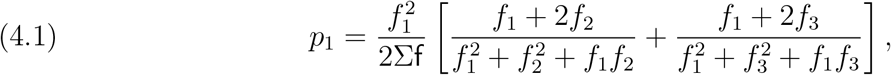

because the case where the process goes through associates of (*f*_1_, *f*_2_, *f*_1_) is symmetrical. Using symmetry again, we obtain

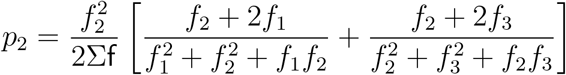

and

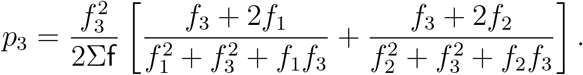

As remarked above, these formulae illustrate the fact that the drift and selection process, besides striving to minimize the number of variants, tries also to maximize the total fitness of the population by selecting against the variants with small fitness.

Analogous formulae for the case *N* = 4 were also obtained, using Maple, but even after simplification, they were too long to be informative; each of them occupied half a page. Hence, in the absence of explicit formulae, we content ourselves with the following theorem which shows that drift and selection events ‘on average’ increase the total fitness of population.

### Theorem 4.3.

*Let* f′ *be the state of the process right after drift and selection event of a population* f. *Then*

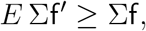

*where E denotes expected value*.

*Proof*. Each event of replacing the *i*th coordinate of f by its *j*th coordinate is paired by the event in which the *j*th coordinate is replaced by the *i*th coordinate. The first of these events takes place with probability 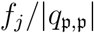, where 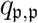 is the diagonal element of the generator matrix in Equ. (3.6). Accordingly, Σf′ – Σf = *f_j_* – *f_i_*, and the second event’s characteristics are symmetrical. Therefore, *E* Σf′ – Σf equals

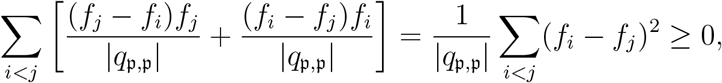

completing the proof.

Next, we turn our attention to the situation where in a homogeneous population a new, possibly better fitted, variant shows up. In other words, the vector of fitnesses is of the form (*f*_1_, *f*_2_, *f*_2_,…, *f*_2_) where *f*_1_ ≫ *f*_2_. We are interested in the probability that variant with fitness *f*_1_ will take over the entire population, i.e. in the probability that the drift and selection process will be absorbed in the state (*f*_1_, *f*_1_,…, *f*_1_).

### Theorem 4.4.

*Let N* ≥ 2. *The probability that the drift and selection process starting at* 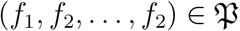 *is eventually absorbed at* (*f*_1_, *f*_1_,…, *f*_1_) *equals*

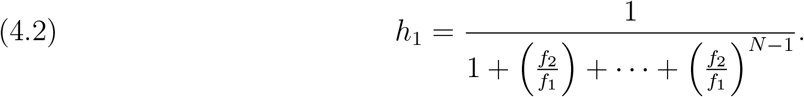

*Proof*. Let *X*(*t*) be the number of individuals of fitness *f*_1_ in the population 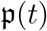. Because of Theorem 4.2, *X*(*t*), *t* ≥ 0 is a time-continuous Markov chain with values in the set 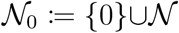, and the elements of its intensity matrix are as follows:

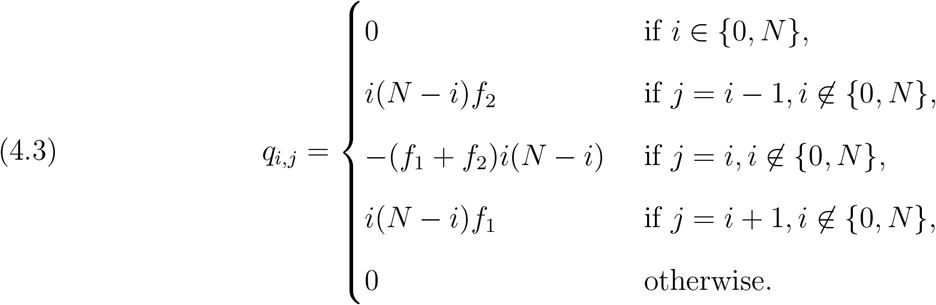

We are interested in the probability that *X*(*t*),*t* ≥ 0 will be absorbed at the state *N*, given that *X*(0) = 1. This probability is the second coordinate in the vector (*h*_0_,…, *h_N_*) of so-called hitting probabilities for the absorbing state {*N*} which, by Theorem 3.3.1 in [25] satisfy the following system of equations:

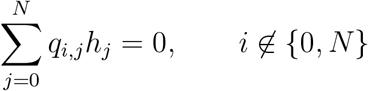

with ‘boundary conditions’ *h*_0_ = 0, *h_N_* = 1. In other words

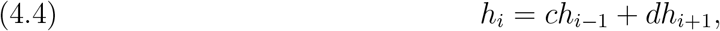

where 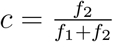 and 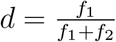 for *i* = 1,…, *N* – 1.

To solve this system, as in p. 16 of [25] or [11] p. 192, we introduce *u_i_* = *h*_*i*−1_ – *h_i_* for *i* = 1,…, *N*. Then the recurrence relation (4.4) becomes 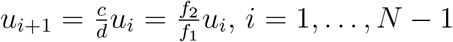.

Therefore, by induction,

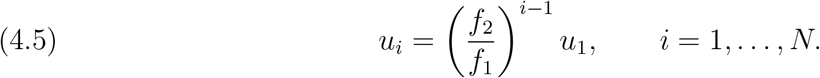

It follows that 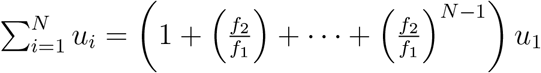. On the other hand, by the definition of *u_i_*s and the boundary conditions for *h_i_*s, 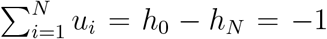. Hence we obtain that 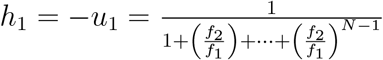, as desired.

We complement this theorem with three remarks. First, we note that (4.5) hides an explicit formula for *h_i_* for any 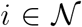, i.e. for the probability that a subpopulation of *i* individuals of fitness *f*_1_ will take over the entire population. For, since 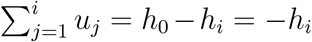, this formula, when combined with (4.2), renders

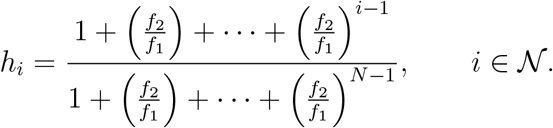

Second, observe that formula (4.2) in the case *N* = 2 agrees with the formula for *p*_1_, and in the case *N* = 3 can be obtained from (4.1) by replacing *f*_3_ by *f*_2_.

Third, in our proof of (4.2) we never used the assumption that *f*_1_ is larger than *f*_2_. However, since *N*, being the size of the considered population, is typically rather large, for *f*_1_ ≤ *f*_2_, the probability of (4.2) is small. This means that new variants without sufficient selective advantage are simply washed away from the population.

On the other hand, given *r* ∈ (0,1) (to play the role of a probability), think of *f*_1_ as of chosen so large as compared to *f*_2_ that

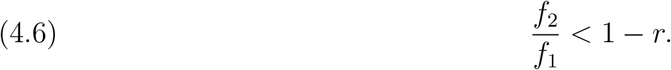

Then 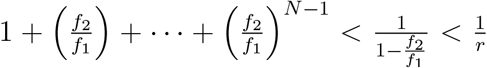. Therefore, for such *f*_1_ (and *f*_2_), *h*_1_ > *r*. In other words, by enlarging *f*_1_ sufficiently, we may make the probability of mutant’s fixation as large as we wish.

We complete this section with information on the expected time to allele’s fixation.

### Theorem 4.5.

*Let N* ≥ 2. *The expected time for the drift and selection process starting at* 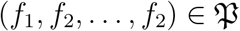 *to be eventually absorbed at* (*f*_1_, *f*_1_,…, *f*_1_) *or* (*f*_2_, *f*_2_,…, *f*_2_) *equals*

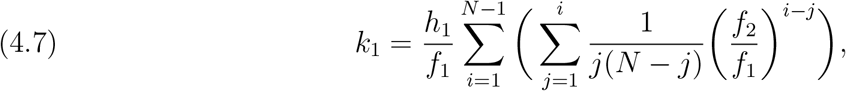

*where h*_1_ *is defined by formula* (4.2).

*Proof*. The time of interest is the second coordinate in the vector (*k*_0_,…, *k_N_*) of hitting times for the absorbing set {0, *N*} which, by Theorem 3.3.3 in [25] satisfy the following system of equations:

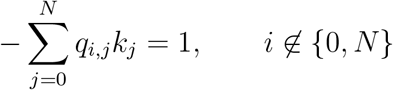

with ‘boundary conditions’ *k*_0_ = 0, *k_N_* = 0, where *q_i,j_*’s are defined in (4.3). In other words

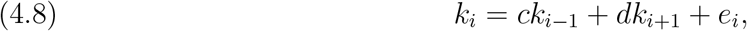

where 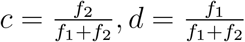 and 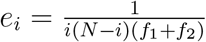 for *i* = 1,…, *N* – 1.

To solve this system, as in the previous theorem, we introduce *v_i_* = *k_i–1_* – *k_i_* for *i* = 1,…, *N*.

Then the recurrence relation (4.8) becomes 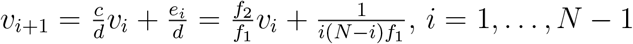. It follows that

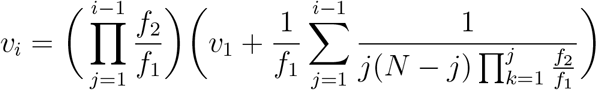

for *i* = 2,…, *N*. We see that

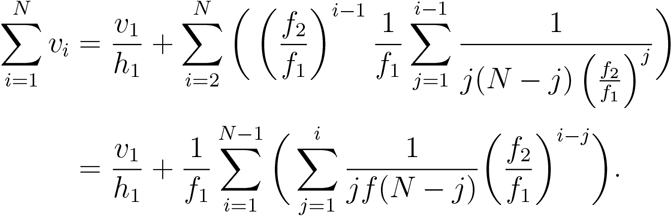

On the other hand, by the definition of *v_i_*’s and the boundary conditions for *k_i_*’s, 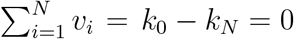. Hence

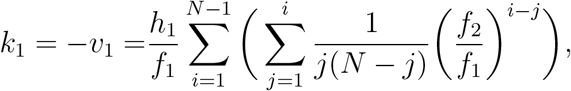

completing the proof.

## 5. Asymptotic behavior of {*P_S_*(*t*), *t* ≥ 0}

In the Supplement 10, we demonstrated the existence of a Markov chain related to the intensity matrix *Q* = *Q_M_* + *Q_S_* of (3.7), i.e. the chain encompassing the mutation and drift “components” of the model we consider.

Before embarking on the study of limit versions of the semigroup {*P*(*t*),*t* ≥ 0} related to this chain, let us rephrase the results of Section 4 to find the limit

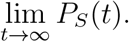

To this end, given 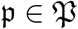, consider

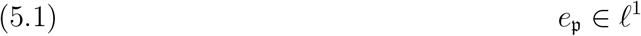

defined by 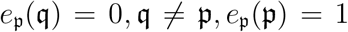. Then 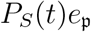 is the 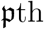 row of transition probability matrix *P_S_*(*t*), composed of probabilities 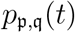 that drift and selection chain starting at 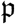 will be at 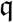 at time *t* ≥ 0. As explained in Section 4 at most *N^N^* probabilities in this row are non-zero, and as *t* → ∞ even all of these at most *N_N_* probabilities tend to zero, save for *M* ≤ *N* of them corresponding to populations where all individuals are identical to each other and to one of the members of the original population, where 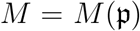 is the number of variants in the population 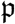. Each of the latter *M* probabilities, on the other hand, converges to one of the probabilities 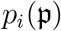 described in Theorem 4.1. Since 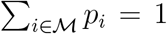, by Scheffés’ Theorem (see e.g. [4]), 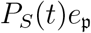 converges to the vector with probabilities 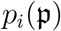, in the norm of *ℓ*^1^. Here is a consequence of this remark.

### Theorem 5.1.

*Let the matrix* 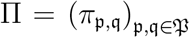 *be defined as follows. For each* 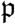 *we choose a subset* 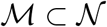 *as described in Theorem 4.1, and let*

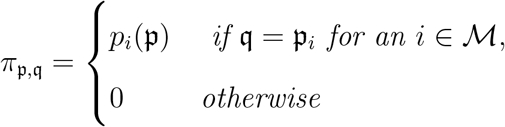

(*where* 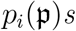 *are defined in Theorem 4.1). Then, for any x* ∈ *ℓ*^1^,

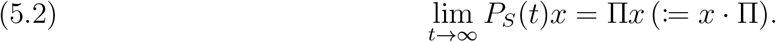

*Proof*. By the reasoning presented above, (5.2) holds for 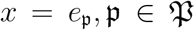. By linearity, this formula extends to all combinations of 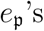. Since such combinations are dense in *ℓ*^1^, a three epsilon argument based on ||*P_S_*(*t*)|| = 1, *t* ≥ 0 completes the proof.

## 6. Asymptotic behavior of {*P*(*t*), *t* ≥ 0}: intuitions

The semigroup {*P*(*t*), *t* ≥ 0} describes the chain in which both mutations and drift and selection events take place. As such, it describes not only a tug-of-war between driver and passenger mutations but also a competition between selections and mutations, these population genetic forces counteracting each other. But, it is one of the main characteristics of drift and selection chain (see (3.4) and (3.6)) that the rate at which new drift and selection events come about grows with the total fitness of the population. It follows that in some regions of 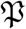 the rate of mutation is larger than the rate of drift and selection events and in other regions the former is smaller than the latter. Hence, in some regions selection will be more expressed, and in other regions effects of mutations will be more apparent.

There are no clear boundaries between these regions, no man’s lands lie between them, and random forces may lead via these no man’s lands from one region to another. Nevertheless, the three main regions, denoted *R*_0_, *R_l_* and *R_u_*, may be characterized as follows.

### 6.1. *R*_0_ region

The central region *R*_0_ contains populations in which drift and selection events occur at a rate that is of the same order as the rate of mutations. By suitable scaling of parameters, this is the region where

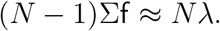

The expression above is inaccurate, since on the left-hand side, instead of the exact expression for the intensity of time to the drift and selection event, we placed a simplified one, true only when all individuals are different. However, this is sufficient for the present purposes which is to define a region in which mutations have force comparable to drift and selection. We will carry out a more accurate analysis using the limit process (see Lemma 9.1 and the text preceding and following it).

An individual member of a population in this region collects new driver and passenger mutations over time: being characterized initially, at time *t* = 0, by (*α, β*), by the time *t* > 0 it becomes of the type (*α* + *m, β* + *n*) (provided it is still alive), but if assumption (2.3) is satisfied, the quotient fitness 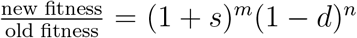 is roughly 1. In other words, between drift and selection events, all individuals travel the path where 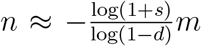; individual fitness does not change much in time. Travels along such paths are of course interrupted by deaths of individuals which are replaced by copies of other individuals. Genetic drift may thus sweep away rare variants, but some kind of statistical equilibrium is obtained between mutations introducing new variants and drift that continually reduces variability (see e.g. [5, 6, 7, 8]).

However, random fluctuations may force a population out of this region of balanced genetic forces to one of the following two regions, where one force prevails against the other. A population may, even the more, be forced out of this region by a temporary or permanent change of parameters *a, d* and *p*, so that e.g. condition (2.4) rather than (2.3) is satisfied.

### 6.2. *R_l_* region

In the lower region *R_l_* we have

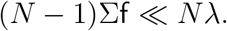

Because of that assumption, drift and selection events are very seldom as compared to mutation events. This means that individuals live for relatively long times, and over periods of their lives accumulate mutations that distinguish them more and more from other individuals. In other words, the members of the population are rather loosely linked, and evolve quite independently of each other.

### 6.3. *R_u_* region

The upper region *R_u_* is characterized by

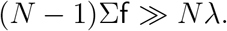

Here the situation is quite different: these are the mutations that are relatively rare as compared to the drift and selection events. Each individual lives for a short time and before it is able to collect a significant number of mutations distinguishing it from other individuals, dies and is replaced by another individual. As a result, very quickly the population becomes uniform: there is practically only one variant in it (i.e., one variant is fixed) as in Theorems 4.1 and 5.1, members of this population could be descendants of a rare but strong variant or of a week but frequent one in the initial population (see Theorem 4.4). In the next two sections we will be able to say more on how mutation process looks like in such populations.

## 7. Asymptotic behavior of {*P*(*t*), *t* ≥ 0} in the upper and lower regions

In this section, we provide a more rigorous mathematical argument, based on the theory of convergence of semigroups, for the intuitions of Sections 6.2 and 6.3. However, this argument still needs to be preceded by the following heuristic reasoning.

Let us consider a subregion *S* of *R_u_* where total fitness of populations, in addition to being much larger’ than *N*λ, is approximately constant: there is a *κ* > 0 such that

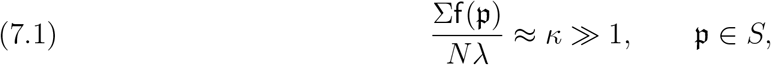

where 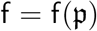 is the fitness vector for 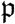. Assume also that for each 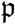 in *S* with the fitness vector f = (*f*_1_,…, *f_N_*) one may find a 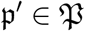 with fitness vector 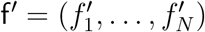 where 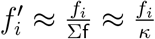. Then 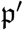 belongs to *R*_0_ and the intensities 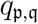 of the drift and selection chain (see (3.4) and (3.6)) in the region S are related to the intensities 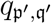 of the corresponding points 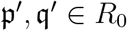 as follows:

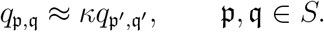

At the same time, intensities of the mutation chain (see (3.3)) do not change in the transfer from 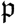 to 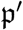.

It follows that instead of thinking of the chain in *S* governed by 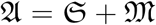 we may think of the chain in *R*_0_ governed by

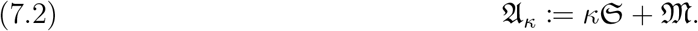

Arguing as at the end of Section 10, we check that for each *κ* > 0, 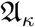 is the generator of a semigroup, say {*P_κ_*(*t*), *t* ≥ 0}, of Markov operators. Thus, our task is that of characterizing the limit

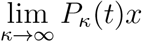

where *x* ∈ *ℓ*^1^ is a distribution concentrated in *R*_0_.

This can be done effectively via Kurtz’s singular perturbation theorem [12, 19, 20] or Chapter 42 in [3], and the analysis does not require assuming that *x* is concentrated in *R*_0_. In a simple case needed in our situation Kurtz’s theorem says that the limit above exists for all *x* ∈ *ℓ*^1^ and *t* > 0 provided the following two conditions are met:

i. lim_*t*→∞_ *P_S_*(*t*)*x* =: Π*x* exists for all *x* ∈ *ℓ*^1^.
ii. 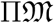 with domain equal to 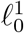 is a generator in 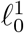, where 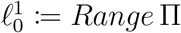.

To deduce this statement from Theorem 42.2 in [3] note that for 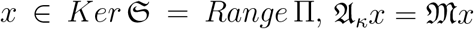 and that for 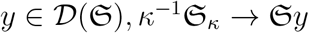. As proved in Theorem 5.1, the first of these two conditions is satisfied, and the second is clear since 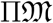 is bounded; this establishes the desired convergence. Moreover, Kurtz’s theorem states that in such a case,

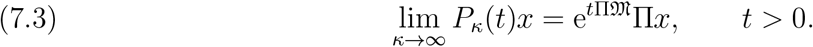

An analogous reasoning shows that instead of thinking about the chain generated by 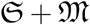 in the lower region *R_l_*, one may think of the chain in *R*_0_ governed by (compare (7.2))

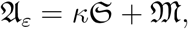

where now *κ* ≪ 1.

The limit

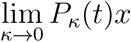

where *x* ∈ *ℓ*^1^ may be found with the help of the Sova-Kurtz version of the Trotter–Kato convergence theorem for semigroups (see [3, 12, 18]): since for each 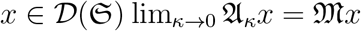, and the set 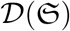 is dense in *l*^1^, we have

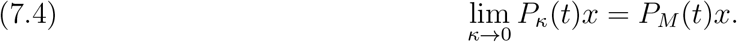

Proofs of the two results are deferred to the supplement. The first one seems to be less intuitive, but we may provide an elementary derivation for the finite dimensional case in which it is sufficient to use Laplace transform and matrix calculus.

### Conjecture

Given matrix exponent

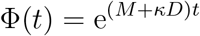

such that

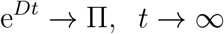

we have

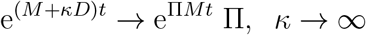

**Proof:** Consider the Laplace transform of Φ(*t*)

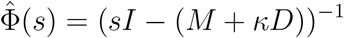

We find that

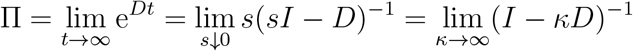

and then

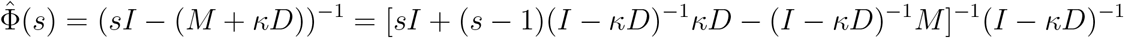

and since

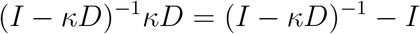

this converges as *κ* → ∞ to

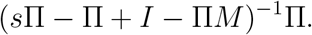

Since Π^2^ = Π the above is equal to (*sI* – Π*M*)^−1^Π, which is the Laplace transform of e^Π*Mt*^ Π, as desired.

## 8. Interpretation of (7.3) and (7.4)

This section is devoted to interpreting the limit theorems just obtained.

### 8.1. Interpretation of (7.3)

Let us start by characterizing the space 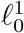 of point (ii) of the previous section. To this end, for *α*, 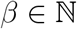, let

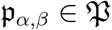

be the population of *N* identical individuals, each of type (*α, β*), and let 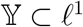 be the subspace spanned by 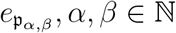 (recall (5.1)). In other words, 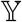 is composed of vectors of the form

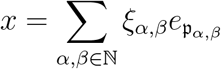

where Σ_*α, β*_ |*ξ_α, β_*| < ∞.

#### Lemma 8.1.

*We have*

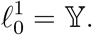

*Proof*. Each 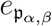 obeys 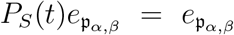, because 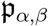 is an absorbing state for the selection/mutation chain. By the definition of Π it follows that 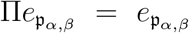, i.e. that 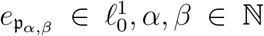. Conversely, in the argument preceding Theorem (5.1) we have shown that for any 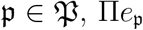 is a convex combination of (a finite number of) vectors 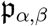, hence is a member of 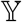. Since 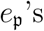 span the entire *ℓ*^1^, this completes the proof.

This lemma shows that the state-space of the limit process (which mimics the selection/mutation/drift process in the regions of high total fitness) is composed of populations in which all individuals are identical: this state-space is

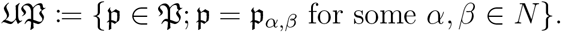

The generator, 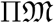, of this process is of interesting form. The value of 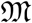 on 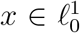 usually does not belong to 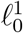. This is because 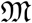 describes mutations: since each and every member of a uniform population may increase the number of driver and/or passenger mutations, after some time the population may contain different variants. However, the generator of the process under consideration is a composition of 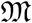 and Π, and the latter operator maps 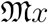 back to 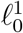. This corresponds to immediate intervening of drift and selection force, which makes the population uniform again, although possibly not quite the same as previously.

Referring back to (3.3) we see 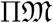 is the generator of a Markov chain in 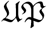 which may be described as follows. Staring at a 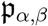 the process stays there for an exponential time with parameter *N*λ. After this time is over, a randomly selected individual (each individual being chosen equally likely) changes its type to (*α* + 1, *β*) with probability *p*, or to (*α, β* + 1) with probability *q*. Then drift and selection either eliminates the new variant or allows it to take over the entire population. According to Theorem 4.4, the variant with new driver mutation is fixed with probability

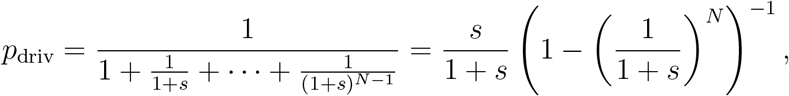

Similarly, the variant with new passenger mutation is fixed with probability

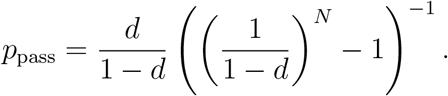

If the new variant is eliminated from the population, everything goes back to the state from before mutation. Thus, the time to effective change is exponential with parameter

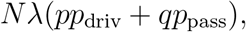

and after this time 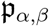 becomes 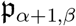 with conditional probability 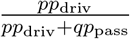 or 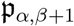 with conditional probability 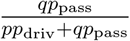.

If 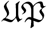 is identified with 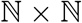, i.e. if each population 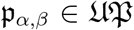 is identified with its type (*α, β*), the process described above is seen to be the pair of two independent Poisson processes on 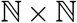: the driver Poisson process with intensity *N*λ*pp*_driv_ (increasing the *α*-coordinate) and the passenger Poisson process with intensity *N*λ*qp*_pass_ (increasing the *β*-coordinate).

Notably, in contrast to the processes of mutations in single individuals, where intensity of driver mutations is much smaller than that of passenger mutations, here the situation is quite the opposite: these are the driver mutations that are typically more frequent than the passenger mutations. For, we have *p*_pass_ ≤ *N*^−1^; on the other hand, arguing as in the vicinity of (4.6), we see that it suffices to take *s* so large that

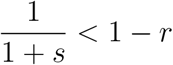

to have

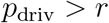

where *r* ∈ (0,1) is given in advance.

The latter phenomenon has its source in the intervening selection process, described above, which eliminates the vast majority of passenger mutations from the population.

Before completing this section, we take a last look at (7.3) and note that this formula informs us also that even though the ‘true’ initial distribution is a member of *ℓ*^1^ and needs not belong to 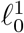, the drift and selection process intervenes so rapidly that before the process of mutations starts the population becomes uniform (this is described by the vector Π*x*). Again, if, for example, in the initial population there is one dominant variant and a single new variant with larger fitness then the latter variant may be fixed with probability given in Theorem 4.4.

### 8.2. Intepretation of (7.4)

Interpretation of (7.4) is much simpler. This formula simply says that in the lower region drift and selection events are so rare that in fact may be disregarded: the chain behaves nearly as if there were no selection or drift. As a result each individual evolves independently of the others, in agreement with intuitions set forth in Section 6.2.

## 9. Simulations

As stated in the Introduction, we consider a population of a fixed number *N* of individuals, each of them characterized by a pair of integers (*α, β*), corresponding to the numbers of drivers and passengers in its genotype, respectively. This pair determines the fitness *f* of the individual by the formula

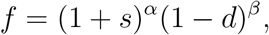

where *s* > 0 and *d* ∈ (0,1) are parameters describing selective advantage of driver mutations over passenger mutations. Thus, the entire population may be identified with the vector

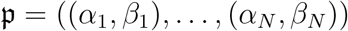

of *N* pairs of integers, with the accompanying vector

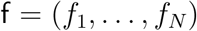

of fitnesses.

In each step of simulation, the decision is made whether the next event is the death and replacement or mutation event. Let us denote by *T_m_* and *T_s_* the exponentially distributed and independent random times to the mutation event and to a death/replacement event, respectively. We simulate both times and the next event occurs at time *t* + min(*T_m_, T_s_*), where *t* is the current time. According to the rules of our process, with mutation rate per cell λ

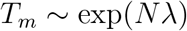

while, following Equ. (3.6)

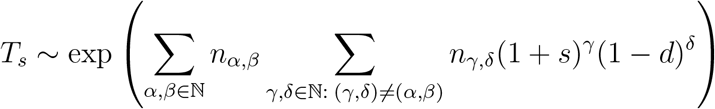

where *n_α,β_* denote the number of individuals of type (*α, β*), so that 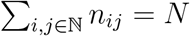.

### Mutation

If *T_m_* < *T_s_*, the next event is mutation. Given this, the index of individual undergoing mutation is drawn from discrete uniform distribution on {1,…, *N*} and the event changes the state of the individual to either (*α*+1, *β*) or (*α*, *β*+1) with probabilities *p* ∈ (0,1) and *q* = 1 – *p* respectively. Fitness of the mutated individual is recalculated accordingly.

### Death and replacement

If *T_m_* ≥ *T_s_*, the next event is death and replacement. Suppose that *K* types of individuals are present, with respective counts *n*(*k*), *k* = 1,…, *K*, summing up to *N*. Following Equ. (3.4), individual *i* with state (*α_i_, β_i_*) is replaced by individual *j* with state (*α_j_, β_j_*), such that (*α_j_, β_j_*) ≠ (*α_i_, β_i_*) with probability proportional to fitness of *j, f_i_* = (1 + *s*)^*α_j_*^ (1 – *d*)^*β_j_*^. The replaced individual inherits the state and fitness from the replacement.

### Trends in fitness

We present a set of stochastic simulations illustrating the richness of possible behaviors of the Tug-of-War process, in its complete and limit versions. One of the issues that we attend to is *criticality*, understood here as the trend of the fitness trajectories, upward, downward or neutral. There are two mathematical facts that provide guidance:

- Mutation vs. selection coefficient balance Equ. (2.4), which indicates the trend in fitness absent drift is determined by the sign of *ps* – *qd*.
- Theorem 4.3, which states that the death and replacement events absent mutation lead to a positive trend in fitness.

Fitness trajectory in the complete process is the result of the interaction of the two trends. If *ps* > *qd*, then the influx of advantageous mutations prevails and in addition, the drift works towards their fixation. As a result the fitness increases rather fast. The effect is subtler when *ps* ≤ *qd*. If the influx of disadvantagous mutations prevails but is not too strong, drift affords to purge the deleterious mutants before they may be fixed. A strong influx is needed to flip this trend.

### Complete process

We follow the interplay between the fluxes *ps* of advantageous and *qd* of deleterious mutations, but also between mutation and drift. The latter can be varied for example by adjusting the parameter *L* = *N*λ. Figure 2 depicts distribution of individual fitness averaged over *N* individuals, in 30 independent runs of the model with a range of parameters. Panel A depicts the case with a *ps* ≫ *qd* (for exact parameter values, see the Figure legend), resulting exponential-like growth of fitness. Panel B shows the case *ps* = *qd*, with the effect being a slow increase of fitness for most runs and a very slow decrease for some. Panel C shows the case of slightly negative trend in mutations *ps* < *qd*. Panel D demonstrates that if *ps* = 0, then drift may efficiently keep purging recurrent deleterious mutants. Panels E and F are showing that in case of increased flux of mutants (large *L*) small changes in the value of *s* parameter, from *s*=0.01 in Fig. 2E to *s*=0.05 in Fig. 2F, may cause a fraction of average fitness trajectories display an upward trend despite large amount of highly deleterious mutations.

**Figure 2.**
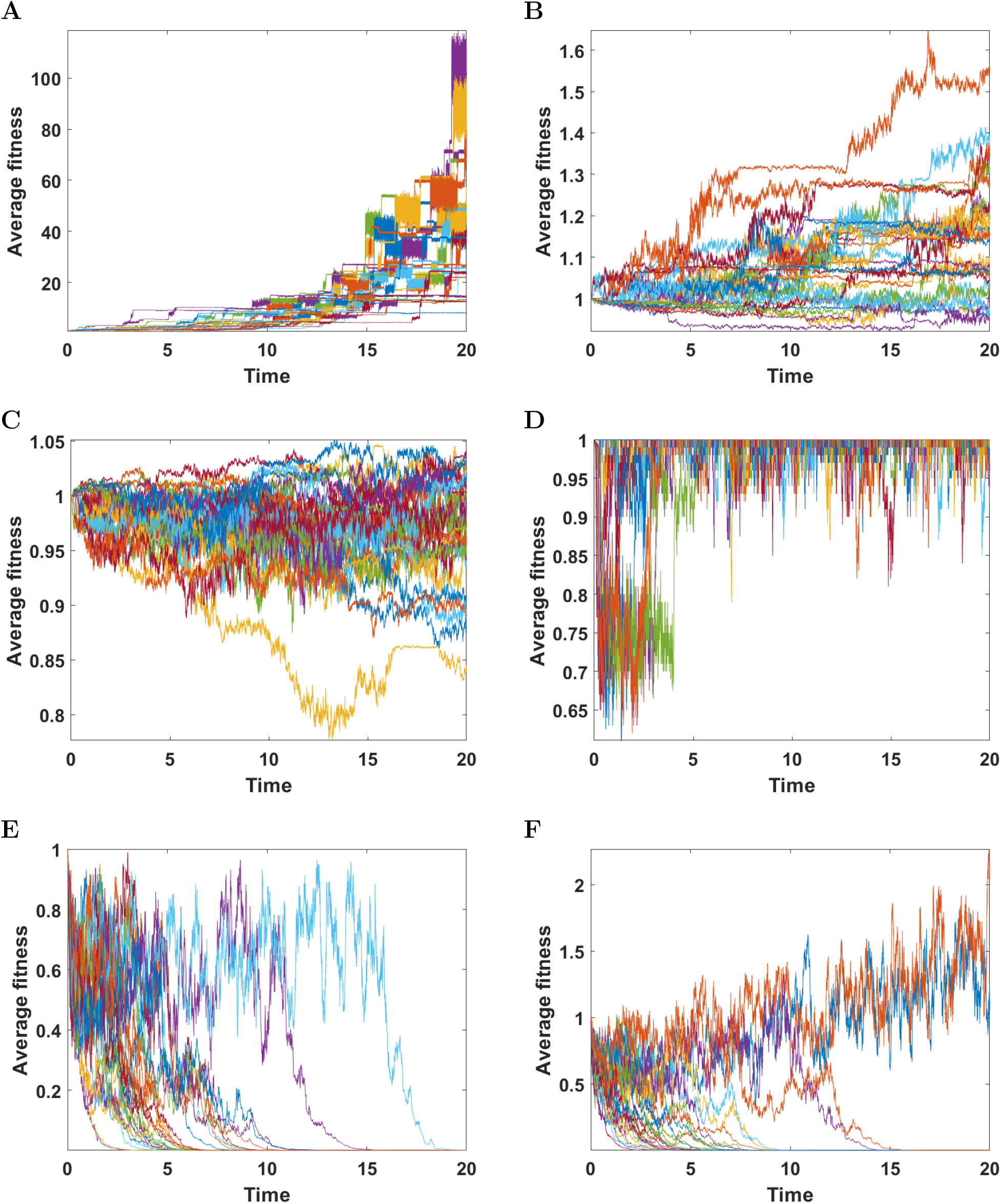
Average fitness of individuals. Results for 30 simulations with parameters: **A**: *s* = 0.8, *d* = 0.05, *p* = 0.1, *L* = 5, *N* = 50 (strong positive selection); **B**: *s* = 0.1, *d* = 0.01111, *p* = 0.1, *L* = 5, *N* = 50 (equilibrium); **C**: *s* = 0.01, *d* = 0.05, *p* = 0.5, *L* = 5, *N* = 50 (negative selection); **D**: *s* = 0, *d* = 0.5, *p* = 0.5, *L* = 5, *N* = 50 (lack of impact of driver mutations); **E**: *s* = 0.01, *d* = 0.5, *p* = 0.1, *L* = 100, *N* = 20; **F**: *s* = 0.05, *d* = 0.5, *p* = 0.1, *L* = 100, *N* = 20 (large mutation rate, passengers prevailing).

Subsequent figures depict runs of a single trajectory of fitness in the process with a range of parameters. Figure 3 depicts one of the average trajectories of Fig. 2A. Panel A depicts the average fitness of population. Mutation events are marked with red (driver) and blue (passenger) asterisks. Let us notice that major jumps in population fitness arise as a result of death-replacement events, more so than of the mutational events, since new arising mutants are frequently purged by death-replacement. Panel B depicts time succession patterns of clones started by driver mutations colored according to fitness of given clone. Panel C depicts genealogies of the clones initiated by drivers and passengers. Lines between nodes represent driver (red) and passenger (blue) mutations. Green circles mark clones alive at the end of simulation (*t* = 20). The vertical axis in panel **C** does not coincide with the time or even with the strict order of clone appearance. However, it is consistent with the ancestor-descendant relationship.

**Figure 3.**
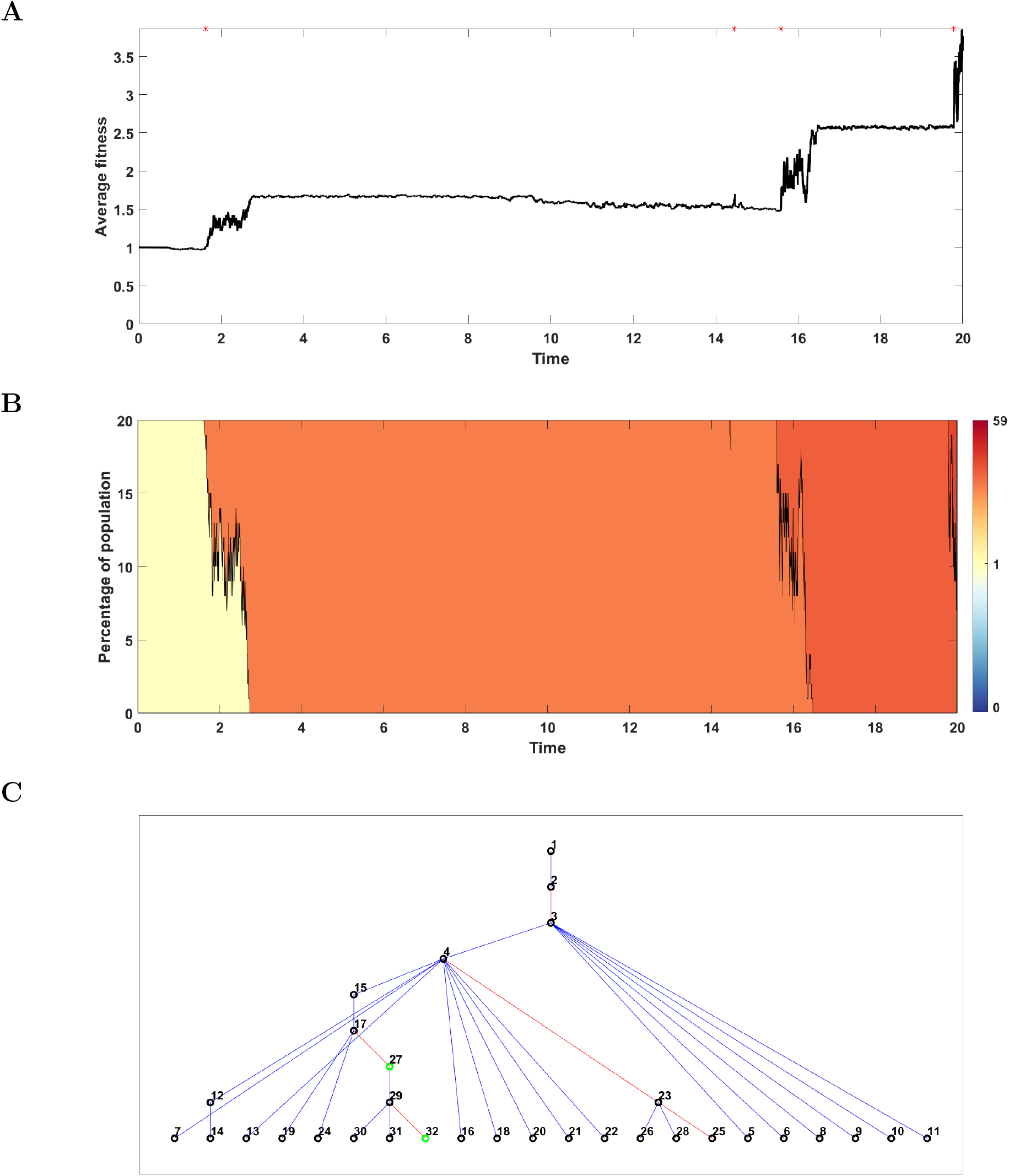
Results for one simulation on *N* = 20 individuals with parameters: *s* = 0.8, *d* = 0.05, *p* = 0.1, *L* = 2 (strong positive selection). **A**: Average fitness of population. Mutation events are marked with red (driver) and blue (passenger) asterisks; **B**: Time succession patterns of clones started with driver mutations colored according to fitness of given clone; **C**: Genealogies of the clones initiated by drivers and passengers. Lines between nodes represent driver (red) and passenger (blue) mutations. Green circles mark are clones alive at the end of simulation (*t* = 20).

Figures 4, 5, 6, 7, 8 and 9 depict single-trajectory plots corresponding all other cases in Fig. 3. Figures corresponding to cases with high mutation rates lack the third panel, since the genealogies of clones become to dense to follow with an increased mutation rate.

**Figure 4.**
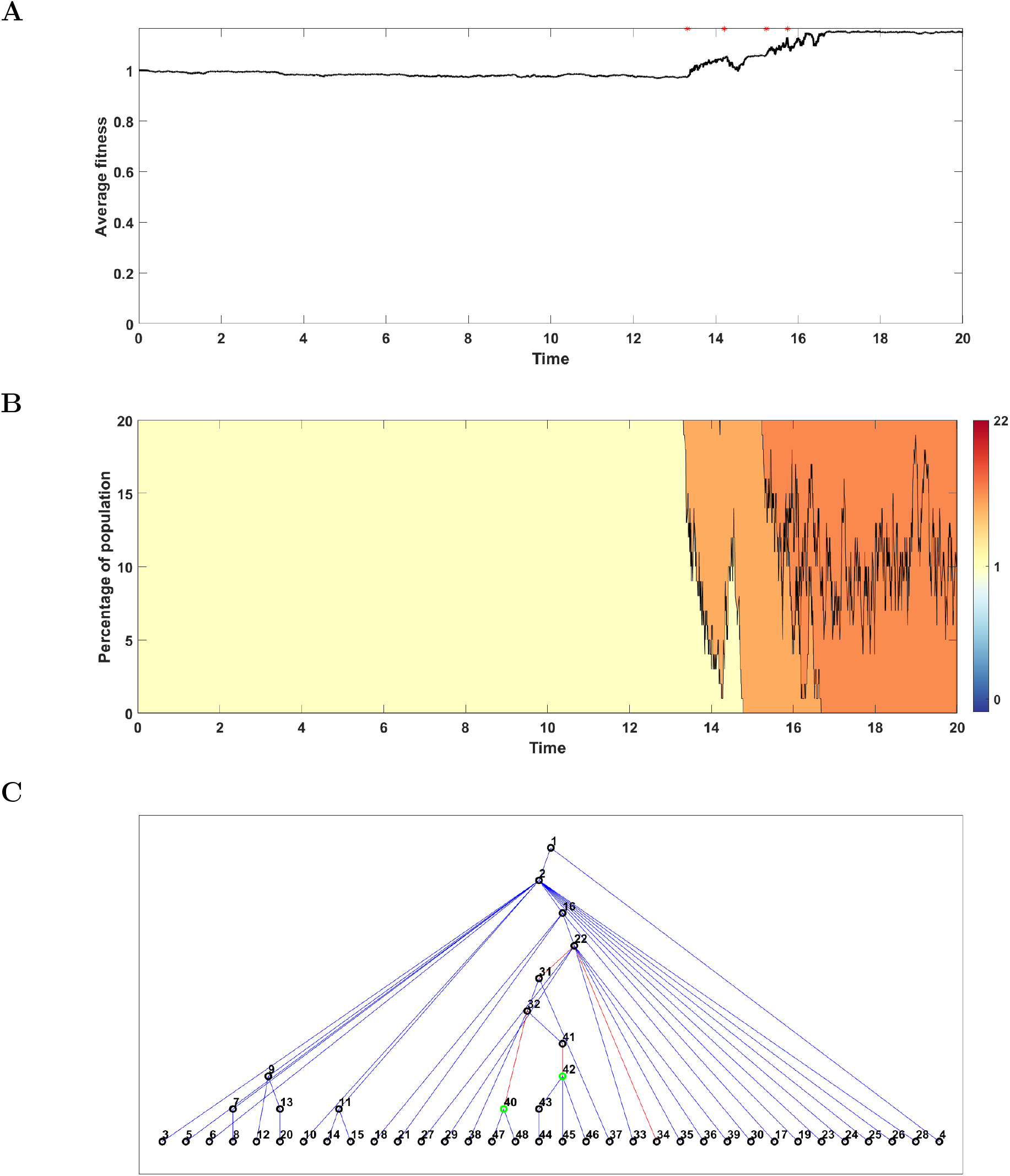
Results for one simulation on *N* = 20 individuals with parameters: *s* = 0.1, *d* = 0.01111, *p* = 0.1, *L* = 2 (equilibrium). **A**: Average fitness of population. Mutation events are marked with red (driver) and blue (passenger) asterisks; **B**: Time succession patterns of clones started with driver mutations colored according to fitness of given clone; **C**: Genealogies of the clones initiated by drivers and passengers. Lines between nodes represent driver (red) and passenger (blue) mutations. Green circles mark clones alive at the end of simulation (*t* = 20).

**Figure 5.**
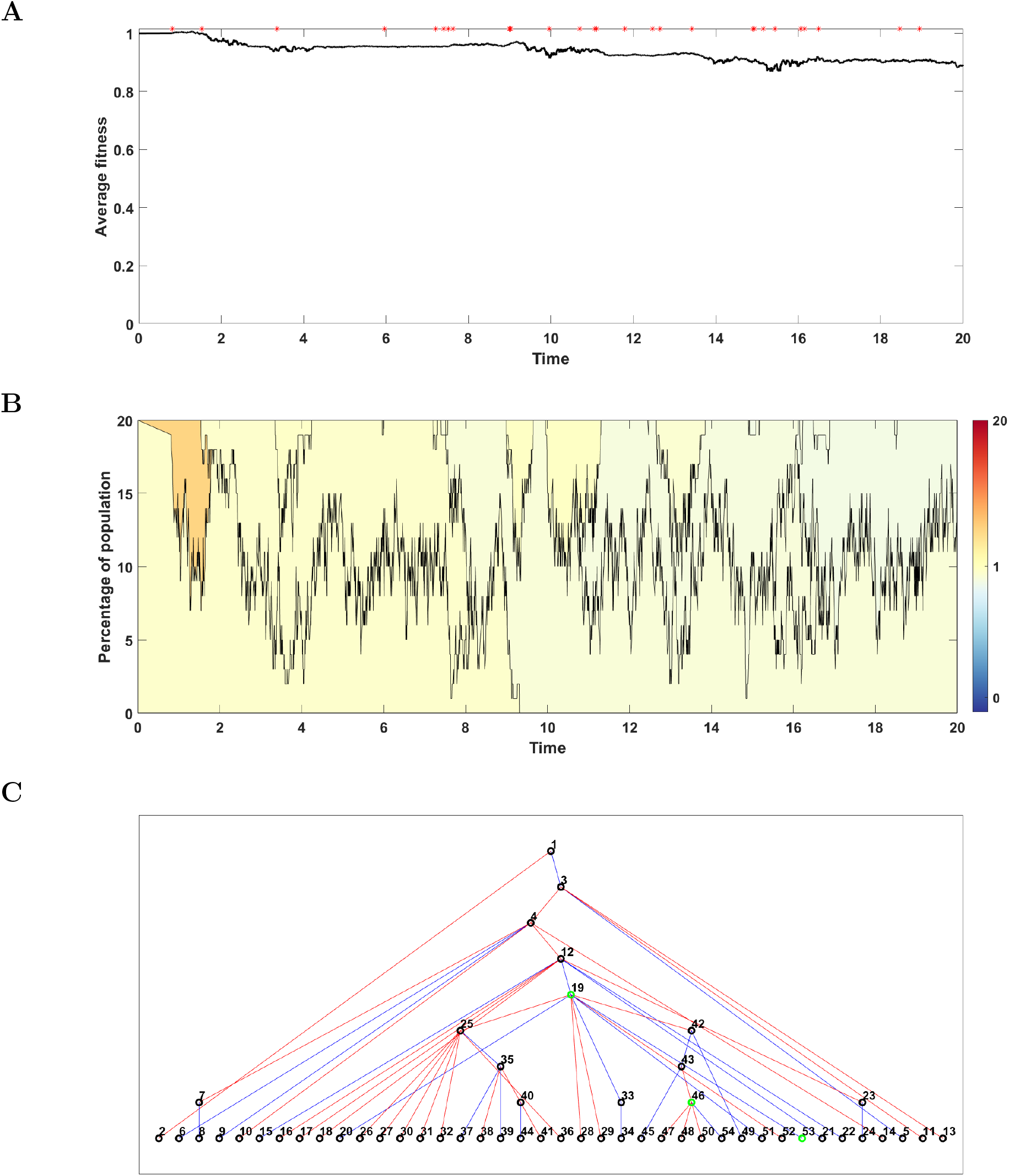
Results for one simulation on *N* = 20 individuals with parameters: *s* = 0.01, *d* = 0.05, *p* = 0.5, *L* = 2 (negative selection). **A**: Average fitness of population. Mutation events are marked with red (driver) and blue (passenger) asterisks; **B**: Time succession patterns of clones started with driver mutations colored according to fitness of given clone; **C**: Genealogies of the clones initiated by drivers and passengers. Lines between nodes represent driver (red) and passenger (blue) mutations. Green circles mark clones alive at the end of simulation (*t* = 20).

**Figure 6.**
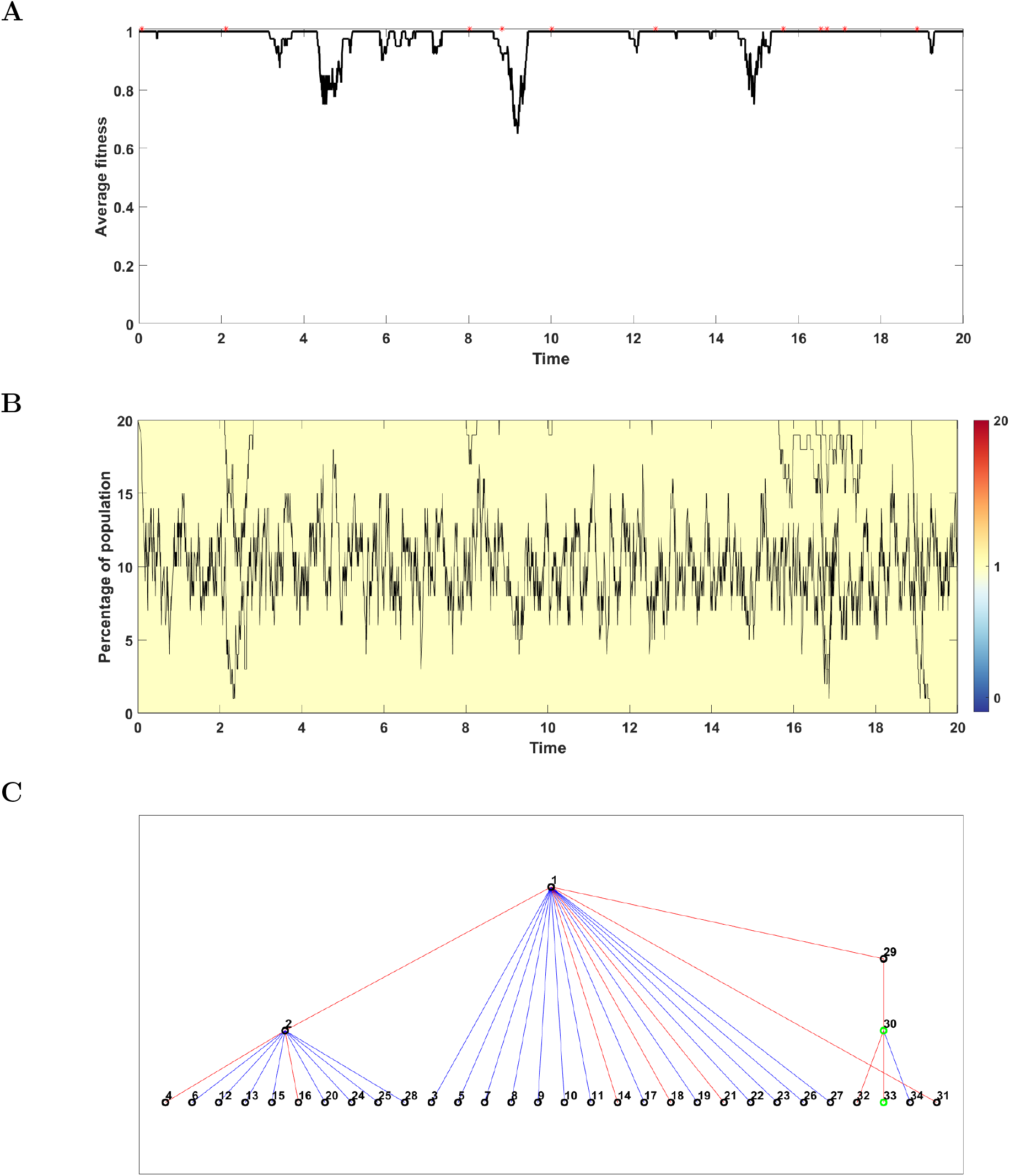
Results for one simulation on *N* = 20 individuals with parameters: *s* = 0, *d* = 0.5, *p* = 0.5, *L* = 2 (lack of impact of driver mutations). **A**: Average fitness of population. Mutation events are marked with red (driver) and blue (passenger) asterisks; **B**: Time succession patterns of clones started with driver mutations colored according to fitness of given clone; **C**: Genealogies of the clones initiated by drivers and passengers. Lines between nodes represent driver (red) and passenger (blue) mutations. Green circles mark clones alive at the end of simulation (*t* = 20).

**Figure 7.**
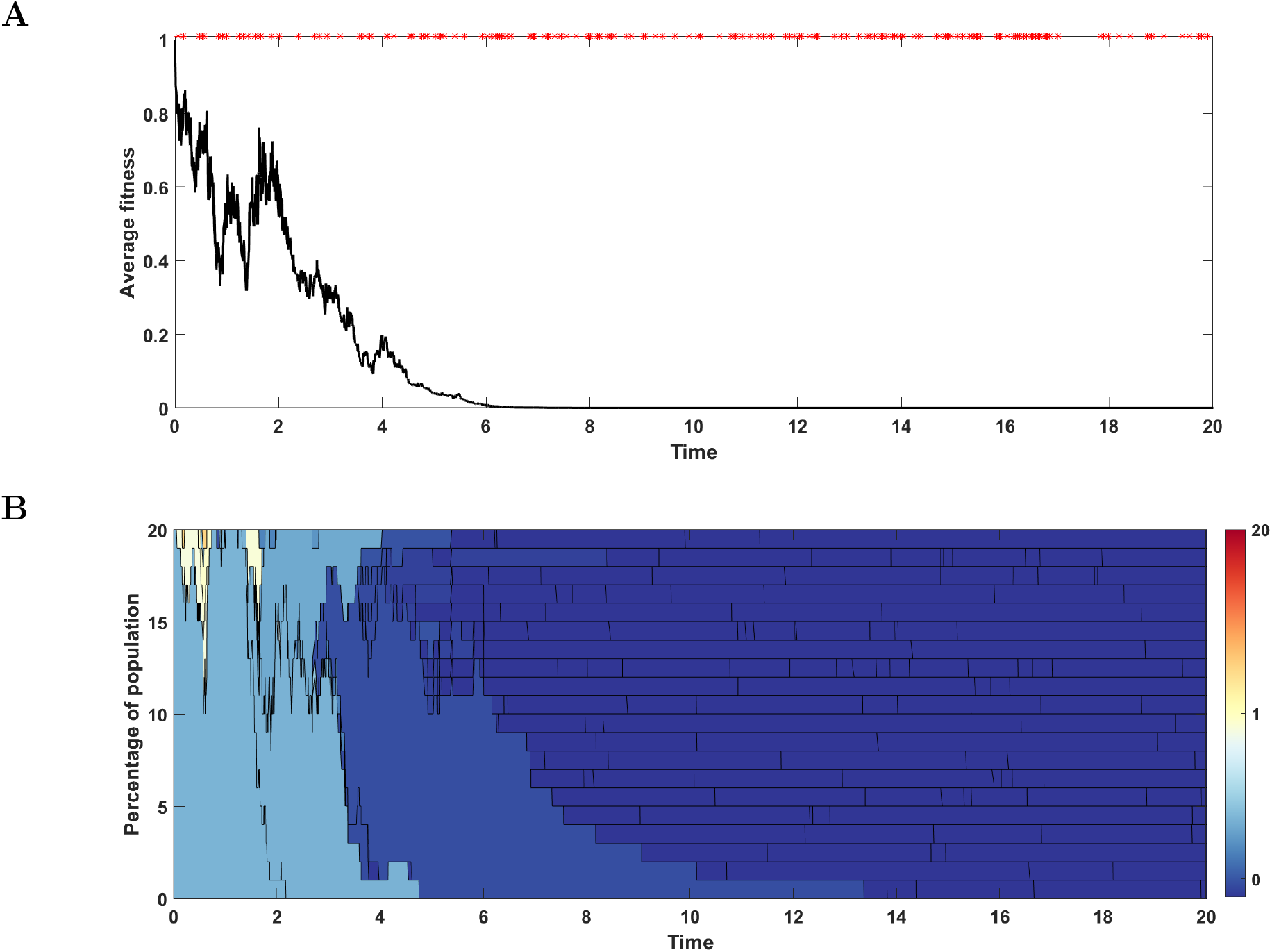
Results for one simulation on *N* = 20 individuals with parameters: *s* = 0.01, *d* = 0.5, *p* = 0.1, *L* = 100 (large mutation rate, passengers prevailing). **A**: Average fitness of population. Driver mutation events are marked with red asterisks; **B**: Time succession patterns of clones started with driver mutations colored according to fitness of given clone.

**Figure 8.**
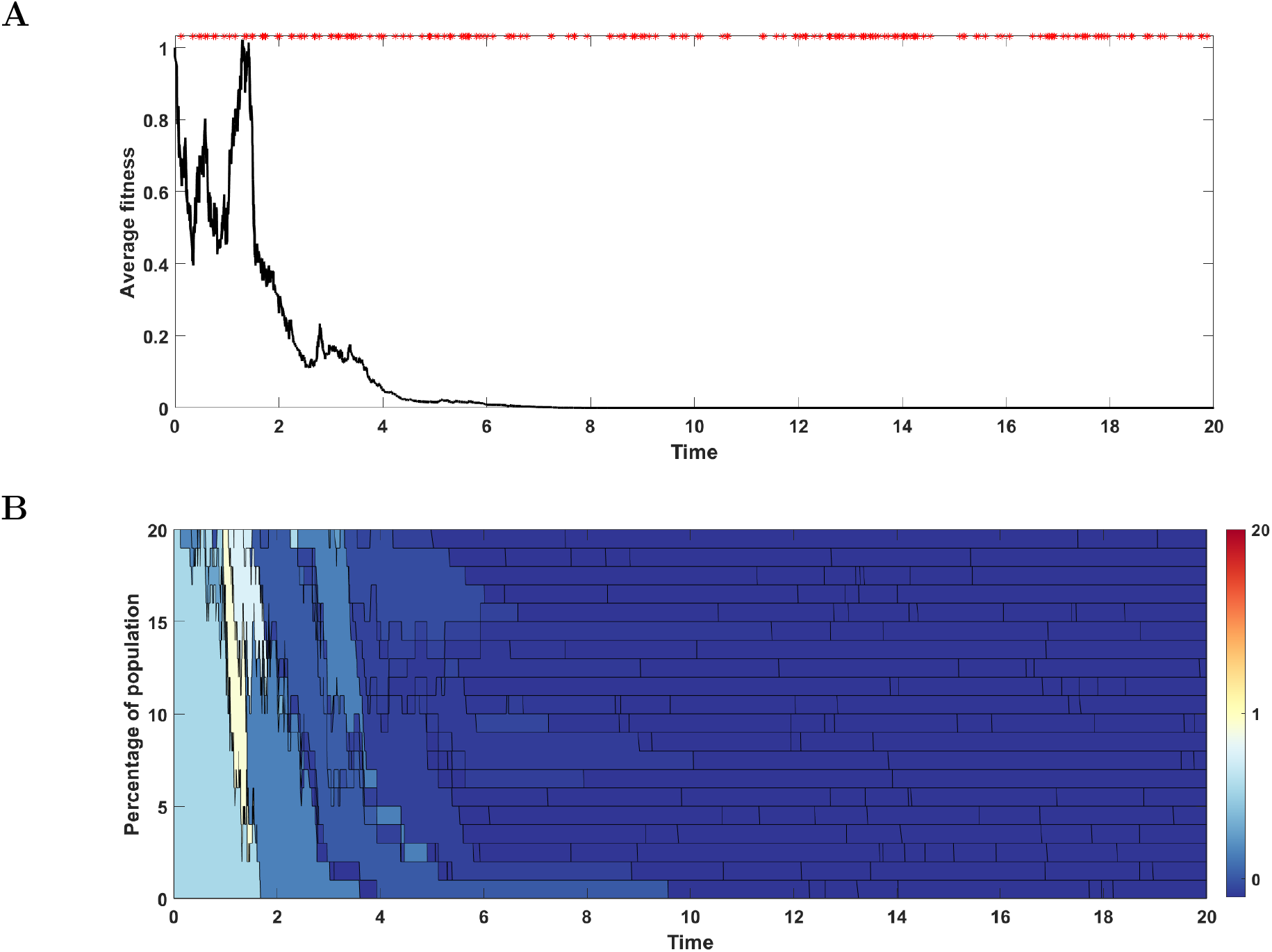
Results for one simulation on *N* = 20 individuals with parameters: *s* = 0.05, *d* = 0.5, *p* = 0.1, *L* = 100 (large mutation rate, passengers prevailing, case with decreasing fitness). **A**: Average fitness of population. Driver mutation events are marked with red asterisks; **B**: Time succession patterns of clones started with driver mutations colored according to fitness of given clone.

**Figure 9.**
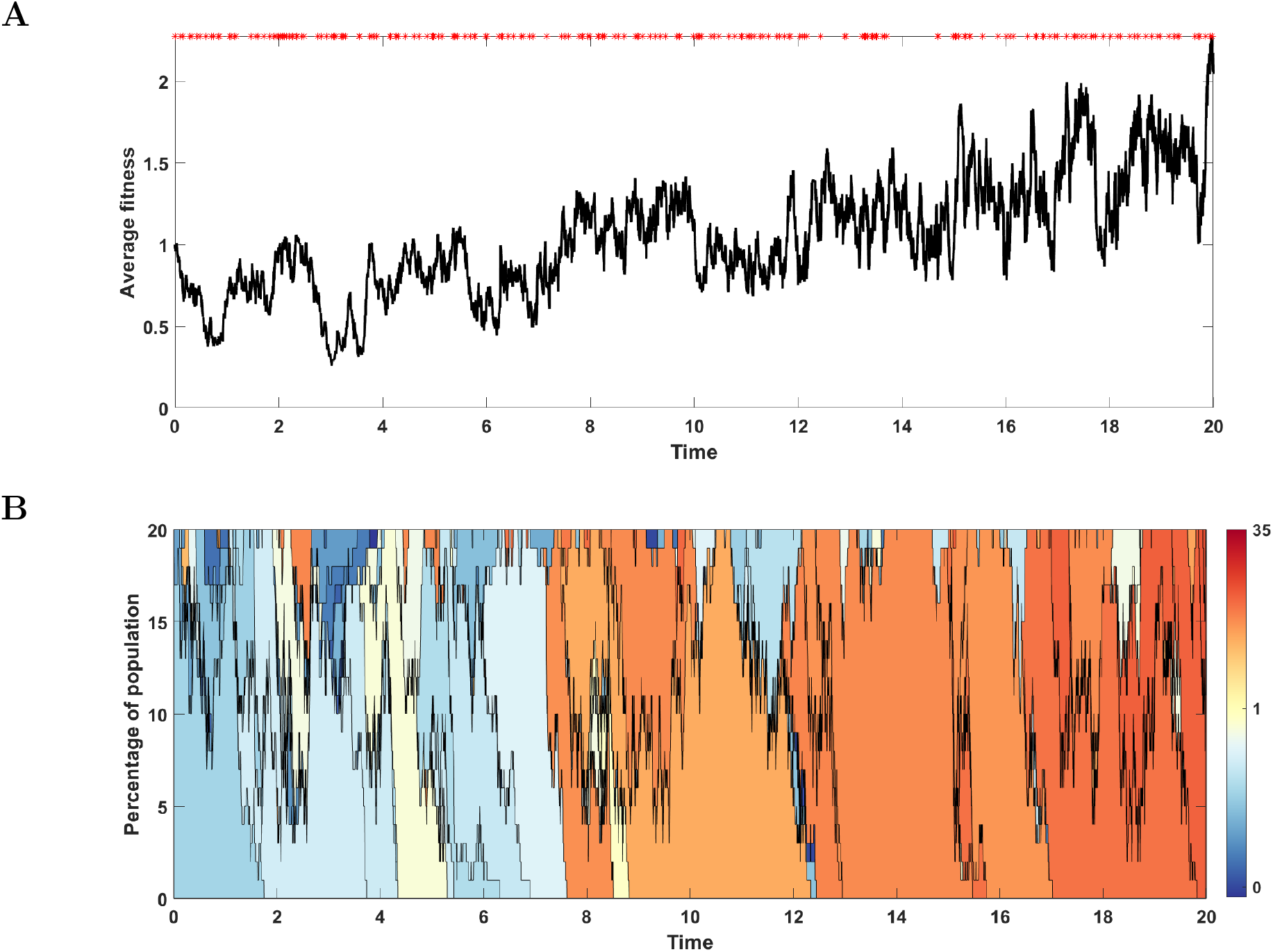
Results for one simulation on *N* = 20 individuals with parameters: *s* = 0.05, *d* = 0.5, *p* = 0.1, *L* = 100 (large mutation rate, passengers prevailing, case with increasing fitness). **A**: Average fitness of population. Driver mutation events are marked with red asterisks; **B**: Time succession patterns of clones started with driver mutations colored according to fitness of given clone.

### Limiting process

Suppose that all individuals have the same fitness (1 + *s*)^*α*^(1 – *d*)^*β*^. The difference between expected fitness right after mutation/drift event and the fitness before this event equals

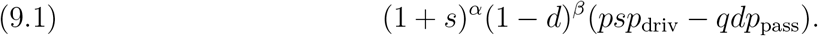

In interpreting this relation we encounter an apparent paradox: in a certain range of parameters, an increase of *d*, that is, a decrease of fitness of passenger mutants, leads to a decrease of the studied difference. In order to explain this paradox we need to consider the function

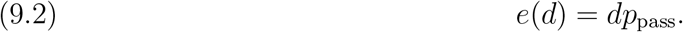

#### Lemma 9.1.

*The function e initially increases and then decreases with d*.

*Proof*. As *d* increases from 0 to 1, 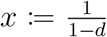 increases from 1 to ∞. Hence, it suffices to check monotonicity of

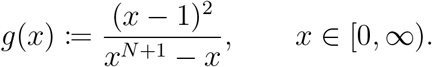

Since

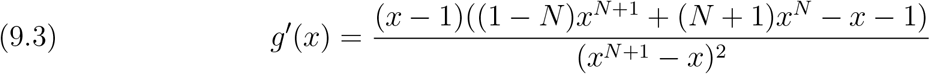

monotonicity of *g* is determined by the sign of

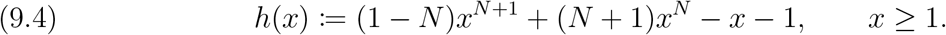

Here *h*(1) = 0 and

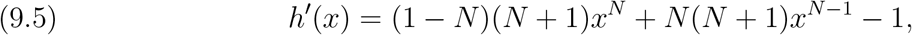

so that in particular *h*′(1) = *N* > 0. Moreover,

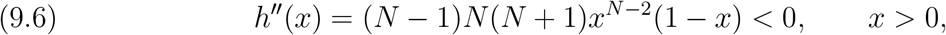

proving that *h*′(*x*) strictly decreases from *N* to −∞ in the interval [1, ∞). Hence, *h* increases from *h*(1) = 0 to a maximum point, and then starts to decrease. Since lim_*x*→∞_ *h*(*x*) = −∞, we conclude that *h* is initially positive, and then, beyond a certain points, say *x*_0_ becomes negative and stays negative for all *x* > *x*_0_. It follows that *g* increases up to *x*_0_ and then starts to decrease, as claimed.

Consider first the scenario in which *e* decreases with the increase of *d*, and thus the difference (9.1) increases. Here, everything seems to agree with our intuition: A decrease in fitness of passenger mutants causes the probability of their fixation to drop and thus if the fitness of driver mutations is the same, the overall population fitness grows faster, because the influence of passenger mutations is smaller.

However, in a certain range of *d*, an increase of *d* (a decrease of fitness of passenger mutations) causes an increase of *e*, and thus a decrease of (9.1). This is because an increase in *d* causes a decrease of *p*_pass_ but this is accompanied by an increase of the first factor in (9.2). It is possible that a change in *d* causes a much smaller change in *p*_pass_ than in *d* itself, and thus may result in the overall growth of *e*. In other words, even though the probability of fixation of a passenger mutation is lower, if such a variant is fixed fitness will drop radically. From this point of view, the observation that a decrease in the fitness of passenger mutants may lead to a decrease of (9.1) is not surprising.

To summarize, analysis of e, the expected drop of the population fitness given that a passenger mutant was fixed, is the key to understanding of the apparent paradox we encountered. It is more informative than the probability *p*_pass_ alone.

The influence of the function

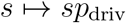

is monotonous; the larger is the fitness of driver mutants, the faster is the growth of the fitness of the entire population.

## 10. Discussion

Following the earlier work of McFarland et al. [21, 22, 23, 24] we build a model of early cancer development which accounts for the influence of two types of mutations: rare drivers with higher fitness, and more frequent passengers with smaller fitness. Mathematically, the model is a continuous-time Markov chain with state-space composed of *N*-tuples of pairs of non-negative integers. Here, *N* is the number of individuals (cells) in the population under study, and is assumed constant; the first coordinate of each pair (individual/cell) is the number of accumulated driver mutations, whereas the second is the number of mutations of passenger type; the resulting individual’s fitness is given by (2.1).

The model may be seen as describing competition of two population genetic forces: selection combined with drift, on one side, and mutations, on the other. Interestingly, the mathematical theory of semigroups of operators, our main tool, allows analysing consequences of these two forces separately, and to infer properties of the full model from the properties of its two components.

The main effect of the first of these components, related to selection combined with drift, is that a population that may initially be heterogeneous, becomes increasingly homogeneous with time. For the associated Markov chain this means that after a random time the process reaches an absorbing state in which all individuals have the same counts of passenger and driver mutations. The corresponding probabilities of fixation of a mutant and the expected times to fixation are calculated in Section 4.

Mutations, on the other hand, introduce new variants to the data at the epochs of a Poisson process; either selectively advantageous drivers, or disadvantageous passengers.

Mathematical analysis of analytical and stochastic properties of the processes related to the two main factors described above allows concluding that they may be combined, and that the new Markov chain that encompasses mutation and selection, on one had, as well as mutations, on the other, is non-explosive. In other words, the underlying stochastic process is a well-defined honest Markov chain.

The resulting process is difficult to analyze. Insights can be obtained using a simpler limit model, presented in Sections 7 and 8, and simulations, see Section 9.

The limit theorem is obtained using the theory of convergence of semigroups of operators [3], and corresponds to the scenario in which the total fitness of the population exceeds certain threshold. The model then predicts that drift and selection events are much more frequent than mutation events. Under such scenario, when a new mutant arises, regardless of whether it is a driver or a passenger, it is almost instantly fixed in the population or completely removed from it. It is the action of the drift and selection chain that causes fixation or removal and favors driver mutations. Therefore, the probability of instantaneous fixation of a passenger mutant is usually smaller than that of a driver mutant. However, because the passengers may arise more often than drivers, the possibility of fixation of passenger mutant is not negligible (see Section 7 for details).

In summary, the limit model state-space is composed of pairs of non-negative integers; this is because each individual is fully characterized by such a pair, and the entire population is composed of identical individuals. At a time a new mutant arises, it is instantly fixed or removed from the population, with probabilities depending on its fitness, and so the population is again homogeneous. Such model seems to account for the influence of driver and passenger mutations. It is interesting that it clearly displays the non-monotonous dependence on the parameter *d* of passenger fitness (Lemma 9.1). Simulations in Figure 10 fully corroborate theoretical analysis.

**Figure 10.**
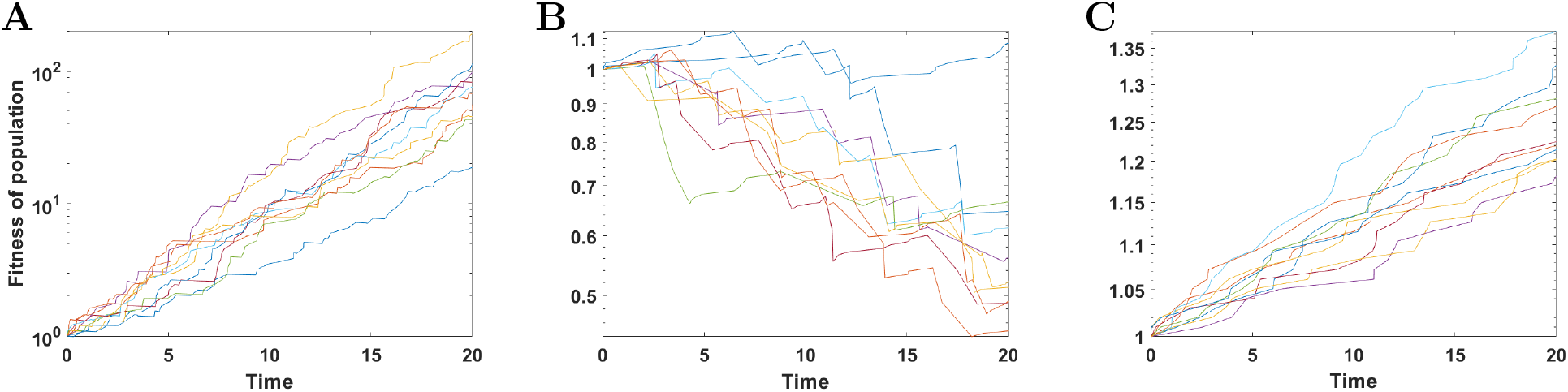
Results for 10 simulations of reduced process with *N* = 20 and remaining parameters: **A**: *s* = 0.1, *d* = 0.01, *p* = 0.5 (*sp* > *dq*); **B**: *s* = 0.01, *d* = 0.1, *p* = 0.5 (*sp* < *dq*); **C**: *s* = 0.01, *d* = 0.5, *p* = 0.5 (*sp* < *dq*). In all cases *L* = 2.)

In addition, simulations show a similar effect in the complete model, as depicted in consecutive panels of Figure 2. In the complete model, the non-monotonicity of the *e*(*d*) corresponds to the balance between downward and upward trends of subsets of trajectories in Figure 2F. The balance is delicate: if the influx of deleterious passenger mutants is limited, drift and selection purge the mutants and population fitness keeps increasing. Only when the influx is sufficiently large, population fitness decreases in part of realization of the process.

We studied by simulation a range of special cases of fitness trajectories and pedigrees of clones originating from driver mutations. A theory of such clones in the Tug-of-war process is still missing. Simulations show how rich is the behavior of this process (Figures 3–9).

With all the reservations, present paper places McFarland’s Tug of War model into the rigorous framework of Moran Model, which allows analyzing it using the well-developed toolbox of time-continuous Markov chains and theory of operator semigroups. Let us notice that our formulation is different from McFarland’s original model as spelled out in [21, 22, 23]. The model there is a state-dependent branching process. To our best knowledge, these models were not rigorously explored. One of the subsequent papers from McFarland’s group [24] explores experimentally the dependence of fitness on the rate of deleterious passenger mutations. However, the references to the mathematical model are only qualitative.

## Supplement: The related semigroup of Markov operators

In Section 4, we have studied in detail the Markov chain related to the intensity matrix *Q_S_*. Since in this chain only a finite number of states can be reached from any given starting point, existence of such chain is an elementary matter. Existence of the chain of mutations (i.e. that related to *Q_M_*) is also clear, as this chain consists of two independent Poisson processes (one for drivers and one for passengers). The question we have never answered is whether there is a Markov chain related to the intensity matrix *Q* of (3.7).

This question is non-trivial because there are so-called *explosive* intensity matrices that are so ‘poorly designed’ that they do not determine the related Markov chain: additional rules need to be specified to describe the chain after the random time of *explosion* (see [4, 9, 25] and references given there). According to the theorem of Kato [16] (discussed e.g. in [1] Chapter 5, [2] pp. 334-338, [3] pp. 74-80, [4] Chapter 3 and [15] pp. 642-647; see also Section 4 in [13] for W. Feller’s proof of this result), for any intensity matrix, whether explosive or not, there is a related minimal Markov chain which, however, is undefined after explosion.

Therefore, in this section we show that *Q* of (3.7) is non-explosive and our argument boils down to the statement that the sum of two intensity matrices, one of which is non-explosive and the other is bounded, is non-explosive. This statement is most naturally proved in the language of semigroups of Markov operators, as we will now explain. Such semigroups are analytical tools for treating Markov chains, and in the later chapters we will use them extensively.

The analysis involves the space 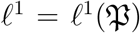 of functions 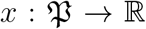 which, because 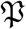 is a countable set, can be considered sequences 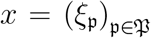 where 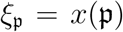 is the value of *x* at 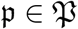. Elements 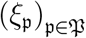 such that 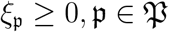 and 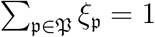 are termed *distributions*.

With each time-continuous Markov chain with values in 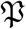 one may associate the probabilities 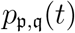 that the chain starting at a 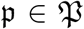 will be at a 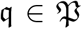 at time *t* ≥ 0. These so-called transition probabilities are conveniently gathered in the matrices

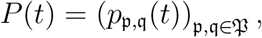

which in turn may be identified (see [4], Chapter 2 for details) with the operators in *ℓ*^1^ defined by the formula

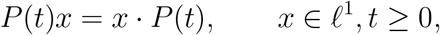

where *x* · *P*(*t*) is the product of a row-vector *x* and the matrix *P*(*t*). All *P*(*t*)’s are Markov operators in that they leave the set of densities invariant: if *x* is an initial distribution of the chain, then *P*(*t*)*x* is its distribution at time *t* ≥ 0. Moreover, the Markov property of the chain is expressed in the semigroup property:

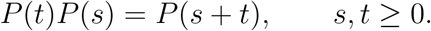

Under mild, natural assumptions on transition probabilities we also have

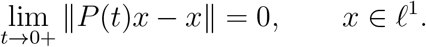

These properties are summarized in the statement that {*P*(*t*), *t* ≥ 0} is a *strongly continuous semigroup of operators* in *ℓ*^1^.

Thus, with each Markov chain we have the associated (uniquely determined) strongly continuous semigroup of Markov operators. Conversely, if all Markov chains with the same transition probabilities are identified, one may speak of *the* Markov chain related to a strongly continuous semigroup of Markov operators.

There are two commonly used infinitesimal descriptions of strongly continuous semigroups of Markov operators in *ℓ*^1^. First, (see e.g. [2, 12, 15]) a strongly continuous semigroup determines and is determined by its generator 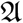, defined by

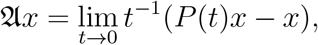

on the domain 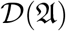 composed of *x* such that the limit on the right-hand side exists. Second, as proved by Doob [10] (see [4, 14]) the limits, called intensities,

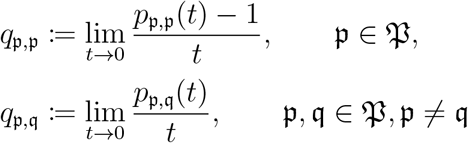

exist, and 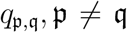 are finite. However, even if all intensities are finite, knowing the entire intensity matrix 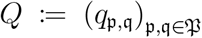 is not equivalent to knowing 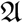. For, whereas 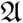 contains the entire information on the semigroup {*P*(*t*), *t* ≥ 0}, the matrix *Q* in general does not. Nevertheless, if *Q* is non-explosive, *Q* and 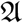 may be somewhat identified: for a typical 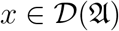, the product *x* · *Q* can be computed, and 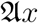 turns out to be equal to this product. For explosive intensity matrices this is not the case; see e.g. the already cited [1, 4], and in particular Chapter 3 in [4]. In fact, for an explosive matrix there are many different Markov chains, many different semigroups of Markov operators and many different generators related to this matrix.

Coming back to the chain of interest, Section 4, and specifically Theorem 4.1 imply that there exists a Markov chain related to the intensity matrix *Q_S_* of (3.4) and (3.6) which is well-defined for all *t* ≥ 0. In particular, *Q_S_* is non-explosive. This is because the related chain reaches an absorbing state by passing through a finite number of transient states. This rules out the possibility of explosion, since an exploding chain is passing through an infinite number of states in a finite time. Therefore, by Kato’ Theorem (see the references earlier on), there is a unique strongly continuous semigroup of Markov operators {*P_S_*(*t*), *t* ≥ 0} in *ℓ*^1^ with the generator, say 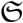, identified with *Q_S_*.

The case of intensity matrix *Q_M_* of (3.3) is simpler, because all its entries are bounded in absolute value by *N*λ, while a bound does not exits for the matrix (3.4)–(3.6). It follows that for *any x* ∈ *ℓ*^1^ the product *x* · *Q_M_* may be computed and belongs to *ℓ*^1^, where *x* is a row-vector, and the map

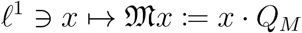

is bounded. Hence, *Q_M_* may be identified with a bounded linear operator 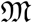 and the semigroup of Markov operators related to *Q_M_* may be defined as the exponent of this operator:

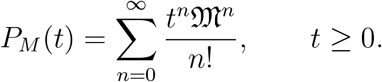

(See e.g. [4] Section 2.3 for details.). Further, operator 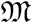 is the generator of semigroup {*P_M_*(*t*), *t* ≥ 0}.

Boundedness of the operator 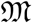 guarantees that the operator

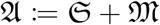

is well-defined on 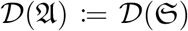 and, in view of the Phillips Perturbation Theorem (see e.g. [2, 4, 12, 15]), is a generator of a strongly continuous semigroup, say {*P*(*t*), *t* ≥ 0}. On the other hand, using the Trotter Product Formula, which says (see the monographs cited above) that

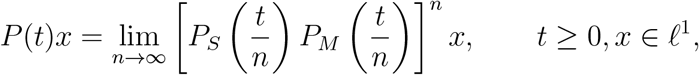

we check that this semigroup is composed of Markov operators, because so are {*P_S_*(*t*), *t* ≥ 0} and {*P_M_*(*t*), *t* ≥ 0}. It can be argued that this semigroup describes the minimal Markov chain related to *Q*. But, by Kato’s Theorem, if *Q* were explosive, this semigroup could not be composed of Markov operators. This shows that *Q* is non-explosive, and thus that the minimal chain is well-defined for all times *t* ≥ 0. It is this minimal chain related to *Q* that models the evolution of our population under selection, drift and mutations. In other words, by *the* Markov chain related to *Q* we mean the unique minimal chain related to this matrix: since *Q* is non-explosive this chain is well-defined for all *t* ≥ 0.

## Data Availability

The authors affirm that all data necessary for confirming the conclusions of the article are present within the article, figures, and tables.

## Acknowledgments

AB was supported by the National Science Centre (Poland) grant 2017/25/B/ST1/01804. MK and MKK were supported by the National Science Centre (Poland) grant 2018/29/B/ST7/02550.

## References

[1] J. Banasiak and L. Arlotti. Perturbations of Positive Semigroups with Applications. Springer, 2006.

[2] A. Bobrowski. Functional Analysis for Probability and Stochastic Processes. An Introduction. Cambridge University Press, Cambridge, 2005.

[3] A. Bobrowski. Convergence of One-parameter Operator Semigroups. In Models of Mathematical Biology and Elsewhere, volume 30 of New Mathematical Monographs. Cambridge University Press, Cambridge, 2016.

[4] A. Bobrowski. Generators of Markov Chains. From a Walk in the Interior to a Dance on the Boundary. Cambridge University Press, Cambridge, 2020.

[5] A. Bobrowski and M. Kimmel. Asymptotic behavior of joint distributions of characteristics of a pair of randomly chosen individuals in discrete-time Fisher-Wright models with mutations and drift. Theor. Popul. Biol., 66(4):355–367, 2004.

[6] A. Bobrowski and M. Kimmel. An Operator Semigroup in Mathematical Genetics. Springer Briefs in Applied Sciences and Technology. Mathematical Methods. Springer, 2015.

[7] A. Bobrowski, M. Kimmel, R. Chakraborty, and O. Arino. A semigroup representation and asymptotic behavior of the Fisher-Wright-Moran coalescent. In “Handbook of Statistics 19: Stochastic Processes: Theory and Methods”, Chapter 8, C.R. Rao and D.N. Shanbhag, eds. Elsevier Science, Amsterdam, 2001.

[8] A. Bobrowski, N. Wang, R. Chakraborty, and M. Kimmel. Non-homogeneous infinite sites model under demographic change: Mathematical description and asymptotic behavior of pairwise distributions. Math. Biosci., 175(2):83–115, 2002.

[9] K. L. Chung. Lectures on boundary theory for Markov chains. With the cooperation of Paul-Andre Meyer. Annals of Mathematics Studies, No. 65. Princeton University Press, Princeton, N.J.; University of Tokyo Press, Tokyo, 1970.

[10] J. L. Doob. Stochastic Processes. Wiley, 1953.

[11] R. Durrett. Probability Models for DNA Sequence Evolution. Springer, New York, Second Edition, 2008.

[12] S. N. Ethier and T. G. Kurtz. Markov Processes. Characterization and Convergence. Wiley, New York, 1986.

[13] W. Feller. On boundaries and lateral conditions for the Kolmogorov differential equations. Ann. of Math. (2), 65:527–570, 1957.

[14] D. Freedman. Markov Chains. Holden-Day, Inc., 1971.

[15] E. Hille and R. S. Phillips. Functional Analysis and Semi-Groups. Amer. Math. Soc. Colloq. Publ. 31. Amer. Math. Soc., Providence, R. I., 1957.

[16] T. Kato. On the semi-groups generated by Kolmogoroff’s differential equations. J. Math. Soc. Japan, 6(1):1–15, 03 1954.

[17] J. F. C. Kingman. Poisson Processes. The Clarendon Press, Oxford University Press, New York, 1993.

[18] T. G. Kurtz. Extensions of Trotter’s operator semigroup approximation theorems. J. Functional Analysis, 3:354–375, 1969.

[19] T. G. Kurtz. A limit theorem for perturbed operator semigroups with applications to random evolutions. J. Functional Analysis, 12:55–67, 1973.

[20] T. G. Kurtz. Applications of an abstract perturbation theorem to ordinary differential equations. Houston J. Math., 3(1):67–82, 1977.

[21] C. D. McFarland. The role of deleterious passengers in cancer. PhD thesis, 2014.

[22] C. D. McFarland, K. S. Korolev, G. V. Kryukov, S. R. Sunyaev, and L. A. Mirny. Impact of deleterious passenger mutations on cancer progression. Proceedings of the National Academy of Sciences, 110(8):2910–2915, 2013.

[23] C. D. McFarland, L. A. Mirny, and K. S. Korolev. Tug-of-war between driver and passenger mutations in cancer and other adaptive processes. Proceedings of the National Academy of Sciences, 111(42):15138–15143, 2014.

[24] C. D. McFarland, J. A. Yaglom, J. W. Wojtkowiak, J. G. Scott, D. L. Morse, M. Y. Sherman, and L. A. Mirny. The damaging effect of passenger mutations on cancer progression. Cancer Research, 77(18):4763–4772, 2017.

[25] J. R. Norris. Markov Chains. Cambridge University Press, Cambridge, 1997.

